# Parallel Rap1>RalGEF>Ral and Ras signals sculpt the *C. elegans* nervous system

**DOI:** 10.1101/2021.02.04.429158

**Authors:** Jacob I. Mardick, Neal R. Rasmussen, Bruce Wightman, David J. Reiner

## Abstract

Ras is the most commonly mutated oncogene in humans and uses three oncogenic effectors: Raf, PI3K, and RalGEF activation of Ral. Understanding the importance of RalGEF>Ral signaling in cancer is hampered by the paucity of knowledge about their function in animal development, particularly in cell movements. We found that mutations that disrupt function of RalGEF or Ral enhance migration phenotypes of mutations in genes with established roles in cell migration. We used as a model the migration of the canal associated neurons (CANs), and validated our results in HSN cell migration, neurite guidance, and general animal locomotion. These functions of RalGEF and Ral are specific to their control of Ral signaling output rather than other published functions of these proteins. In this capacity Ral functions cell autonomously as a permissive developmental signal. In contrast, we observed Ras, the canonical activator of RalGEF>Ral signaling in cancer, to function as an instructive signal. Furthermore, we unexpectedly identified a function for the close Ras relative, Rap1, consistent with activation of RalGEF>Ral. These studies define functions of RalGEF>Ral, Rap1 and Ras signaling in morphogenetic processes that fashion the nervous system. We have also defined a model for studying how small GTPases partner with downstream effectors. Taken together, this analysis defines novel molecules and relationships in signaling networks that control cell movements during development of the nervous system.

## INTRODUCTION

Approximately one-third of human tumors harbor activating mutations in Ras, the most commonly mutated oncogene. Ras is the founding member of the Ras superfamily of small GTPases, membrane-tethered GDP/GTP-cycling switches that are positively regulated by GEFs (<u>g</u>uanine nucleotide <u>e</u>xchange factors) and negatively regulated by GAPs (<u>G</u>TPase <u>a</u>ctivating <u>p</u>roteins; (Reiner and Lundquist, 2018; Wennerberg et al., 2005). Ras plays a central role in cell-cell signaling and thus promotes many facets of tumorigenesis, including uncontrolled proliferation, invasion, metastasis, tumor angiogenesis, survival and anchorage-independent growth. Yet, because Ras itself mostly cannot be targeted pharmacologically (Kim et al., 2020), research into small molecule inhibitors has shifted largely to its oncogenic effectors (Shields et al., 2000). Therefore, study of the effectors of Ras is critical to understanding Ras function in development and disease.

Ras oncogenic signaling is propagated by three oncogenic effectors. Activation of the canonical Raf S/T protein kinase triggers signaling through MEK>ERK MAP kinase cascade. Activation of PI3K lipid kinase, which can also be activated via receptors, triggers activation of the PDK-Akt cascade. The Raf and PI3K cascades are among the most studied and pharmacologically targeted cascades in all of biology (Cox et al., 2014).

The third oncogenic Ras effector, RalGEF activation of Ral, is poorly understood. Ral (<u>Ra</u>s-<u>l</u>ike) is itself a small GTPase that has a core effector-binding loop distinct from that of Ras. Ras binds and activates RalGEF, which in turn activates Ral. Ral is an oncogene (Gentry et al., 2014; Kashatus, 2013), and its GAP is a tumor suppressor (Beel et al., 2020; Gao et al., 2019; Iida et al., 2020; Oeckinghaus et al., 2014; Saito et al., 2013; Uegaki et al., 2019). Of the two mammalian Ral orthologs, RalA is associated with tumor initiation and anchorage-independent growth and RalB is associated with invasion, metastasis and survival (Gentry et al., 2014; Kashatus, 2013).

While Ras is accepted as the primary activator of RalGEF>Ral signaling, work in yeast two-hybrid and *in vitro* systems illustrates that the close paralog of Ras, Rap1 (<u>Ra</u>s <u>p</u>roximal), can also bind RalGEF (Frische et al., 2007). Rap1 shares an identical effector binding loop with Ras. Additionally, observations in *Drosophila* argue that Rap1 can also activate RalGEF>Ral signaling (Carmena et al., 2011; Mirey et al., 2003). Yet technical limitations at the time and essential roles for proteins in flies prevented analysis with endogenous proteins, underscoring the value of the developmentally and anatomically simpler *C. elegans*. Rap1 has also been implicated as a mammalian oncoprotein (Shah et al., 2019), though interpretation is confounded by the demonstrated role of Rap1 in promoting cell-cell-junctions and diluting Ras nanoclusters (Nussinov et al., 2020; Pannekoek et al., 2014), both of which may work to oppose tumorigenesis. In *C. elegans*, we found that Rap1^RAP-1^ functions in the capacity of a weaker form of Ras in Ras^LET-60^-dependent developmental events (Rasmussen et al., 2018).

Like Ras, Ral uses three main oncogenic effectors. One of these, RalBP1, is implicated in the generation of invadipodia (Neel et al., 2012). The other two effectors, Exo84 and Sec5, are both components of the heterooctameric exocyst complex. The exocyst plays essential rolls in exocytosis as well as functioning downstream of the PAR complex in establishment and maintenance of apical-basal polarity in tubulogenesis (Armenti et al., 2014; Wu and Guo, 2015). Since the exocyst likely interacts with hundreds of proteins, finding potential Ral-dependent signaling partners via classic biochemical approaches has proved difficult.

*C. elegans* confers the advantage of simple development and single coding genes for all of the protein molecules in the Ras signaling system named here; mammals typically have two to four ortholog-encoding genes. We have used *C. elegans* genetics with endogenous genes and proteins to investigate *in vivo* mechanisms of Ral function and signaling. During induction of vulval cell fates, a classic system for growth factor signaling and the formation of epithelial tubes (Shin et al., 2018), Ras switches effectors, from Raf to RalGEF>Ral (Zand et al., 2011). Ras>RalGEF>Ral functions as a modulatory signal to promote 2° vulval fate in support of the main signal, Notch. Ras>RalGEF>Ral signals through Exo84 of the exocyst and a CNH domain MAP4 kinase, GCK-2, to activate MLK-1/MAP3 kinase and p38/PMK-1 MAP kinase (Shin et al., 2018). RalGEF is bifunctional and promotes 1° fate in addition to its canonical role in promoting 2° fate through Ral, probably as a scaffold the modulatory PI3 Kinase>PDK>Akt cascade (Shin et al., 2019). Ral also activates TORC1 in parallel to Rheb to control lifespan in *C. elegans* and invasion in mammalian cells (Duong et al., 2020; Martin et al., 2014).

Despite inroads in using *C. elegans* as a system to understand Ral-dependent signal transduction, we have not yet identified a role for Ral in morphogenesis, which might provide a model for human RalB in invasion and metastasis. Consequently, here we investigate the role of RalGEF^RGL-1^>Ral^RAL-1^ signaling in morphogenetic events that sculpt the nervous system of *C. elegans*. We describe that in *C. elegans,* RalGEF^RGL-1^>Ral^RAL-1^ signaling alone is not essential for proper cell migrations or sculpting of the nervous system via control of growth cone migration. However, the use of sensitized backgrounds reveals that loss of RalGEF^RGL-1^>Ral^RAL-1^ exacerbates existing migration and guidance defects, suggesting that RalGEF^RGL-1^>Ral^RAL-1^ contributes to many spatial guidance events as a parallel modulatory signal. We investigate this interaction primarily in the migration of the canal associated neurons (CANs), but further test our hypothesis by observing other migratory events of cells and growth cones. Our results also reveal a qualitative difference of contributions of Ras^LET-60^ and RalGEF^RGL-1^>Ral^RAL-1^ to this process, consistent with Ras^LET-60^ mediating an instructive cue while RalGEF^RGL-1^>Ral^RAL-1^ mediates a permissive cue. Accordingly, we observed that deletion of Rap1 phenocopies disruption of RalGEF> Ral signaling. Our results illustrate that RalGEF>Ral functions broadly in the development of the animal, and provide a platform for further investigation of the mechanisms by which Rap1^RAP-1^>RalGEF^RGL-1^>Ral^RAL-1^ signaling contributes to development and is orchestrated independently of Ras^LET-60^.

## MATERIALS AND METHODS

### *C. elegans* handling and genetics

All strains were derived from the N2 Bristol wild type. Animals were grown with *E. coli* OP50 bacteria on NGM agar plates at 20°C unless stated otherwise. Strains used are listed in Supplementary Table 1. Data analysis was performed using GraphPad Prism software (GraphPad Software Inc., La Jolla, CA).

### Genotyping

PCR primers used are listed in Supplementary Table 2. Animal genotyping PCR (Taq PCR Master Mix, Qiagen) samples were run on either 0.8% or 2.0% agarose gels, depending on band sizes. Control +/+, m/+, and m/m reactions were included with each strain construction and gel. All strains constructed using PCR genotyping were confirmed with a second PCR test before freezing and use in assays.

Triplex PCR with primers DJR614/615/616 (Tm=59°C; 35 cycles) was used to detect *rgl-1(ok1921)*, resulting in 366 bp (wild type) and 233 bp (*ok1921*) product bands. *rgl-1(tm2255)* was detected by primers FSM7/8/9 (Tm=58°C; 35 cycles), resulting in 509 bp (wild type) and 254 bp (*tm2255*) product bands. Putative GEF-dead *rgl-1(gk275304)* and nonsense *rgl-1(gk275305)* alleles were tracked in *trans* to *rgl-1(tm2255).* Point mutation *ral-1(gk628801*rf) was amplified by primers DJR778/779 (Tm=59°C; 40 cycles) to generate a 250 bp band, which was then digested with HpyCH4IV (NEB; 2.5 units added to total reaction volume with NEB buffer 3.1 to 0.5x total, digested at 37°C overnight). Because the *gk628801* point mutation eliminates the HpyCH4IV site, after digestion the mutant allele yields 250 bp band while the wild-type allele yields bands of 122 and 128 bp (Shin et al., 2018). Both *ral-1(re218)* and *ral-1(re160*gf) were detected by TD185/186/187 (Tm=52°C; 35 cycles), resulting in band sizes of 1100 bp (wild type) and 908 bp (CRISPR insert) (Shin et al., 2018). *rlbp-1(tm3665)* was detected by triplex PCR with primers REW102/103/104 (Tm=61°C; 35 cycles), resulting in band sizes of 297 bp (wild type) and 505 bp (*tm3665*). *exoc-8(ok2523)* was detected by triplex PCR with primers REW109/110/111 (Tm=61°C; 35 cycles), resulting in band sizes of 411 bp (wild type) and 270 bp (*ok2523*) (Shin et al., 2018). CAN-expressing putative *ral-1* rescuing insertions *reSi8* and *reSi9* in chromosomal position I: −5.32 were detected by triplex PCR with primers JIM064/065/066 (Tm=60°C; 35 cycles), resulting in band sizes of 517 bp (wild type) and 130 bp. CAN-expressing putative *rgl-1* rescuing insertions *reSi14* and *reSi15* in chromosomal position I: −5.32 were detected by triplex PCR with primers JIM064/066/072 (Tm=60°C; 35 cycles), resulting in band sizes of 517 bp (wild type) and 151 bp.

### Plasmids cloning and transgene generation

Plasmid pJM1 (*P_ceh-23_L_::mKate2::rgl-1*) was generated using Gibson Assembly (NEB) of PCR products. The CAN-specific portion of the *ceh-23* promoter (*P_ceh-23_L_*) was amplified from clone pOH128 (Wenick and Hobert, 2004) using primers JIM032/033. Primers JIM034/035 were used to amplify mKate2 from pNR020 (pBS::AID::mKate2::2xHA::AID) and primers JIM036/037 were used to amplify *rgl-1* from pREW21 (Shin et al., 2019), with added overlapping homology arms for pJM1 Gibson Assembly. Plasmid pJM2 (*P_ceh-23_L_::mKate2::ral-1)* was also generated using Gibson Assembly of PCR products. Primers JIM034/038 were used to amplify mKate2 from pNR020 (pBS::AID::mKate2::2xHA::AID) and primers JIM039/040 were used to amplify *ral-1* from pEP4.1 with added overlapping homology arms for pJM2 Gibson Assembly.

Original injections of pJM1 and pJM2 to generate multi-copy transgenes resulted in silencing of mKate2 expression within a few generations. Consequently, the inserts of pJM1 and pJM2 were cloned into the pCC249 plasmid backbone, typically used for miniMos (de la Cova and Greenwald, 2012; Frokjaer-Jensen et al., 2014). However, for more control over location and copy number, we used this plasmid as a repair template for CRISPR insertion into a safe-harbor site on Chromosome I, located at −5.32, the position in which the transposon ttTi4348 is inserted and is frequently used for MosSCI insertions (Frokjaer-Jensen et al., 2008). Primers JIM048/049 were used to amplify the pCC249 backbone for Gibson Assembly with pJM1 and pJM2 inserts. Primers JIM050/051 were used to amplify pJM1 with added homology arms to generate pJM3 while JIM050/052 were used to amplify pJM2 with added homology arms to generate pJM4. Resulting plasmid inserts were inserted via CRISPR in position I: −5.32. mKate2 expression from these lines was not silenced (see Fig 3A-F). The resulting *reSi8, and reSi14 and reSi15* were crossed into *ral-1(gk628801); otIs33*; *mig-2(gm38*gf) and *otIs33*; *rgl-1(tm2255) mig-2(gm38*gf), respectively, to evaluate CAN-specific rescue.

**Figure 1:**
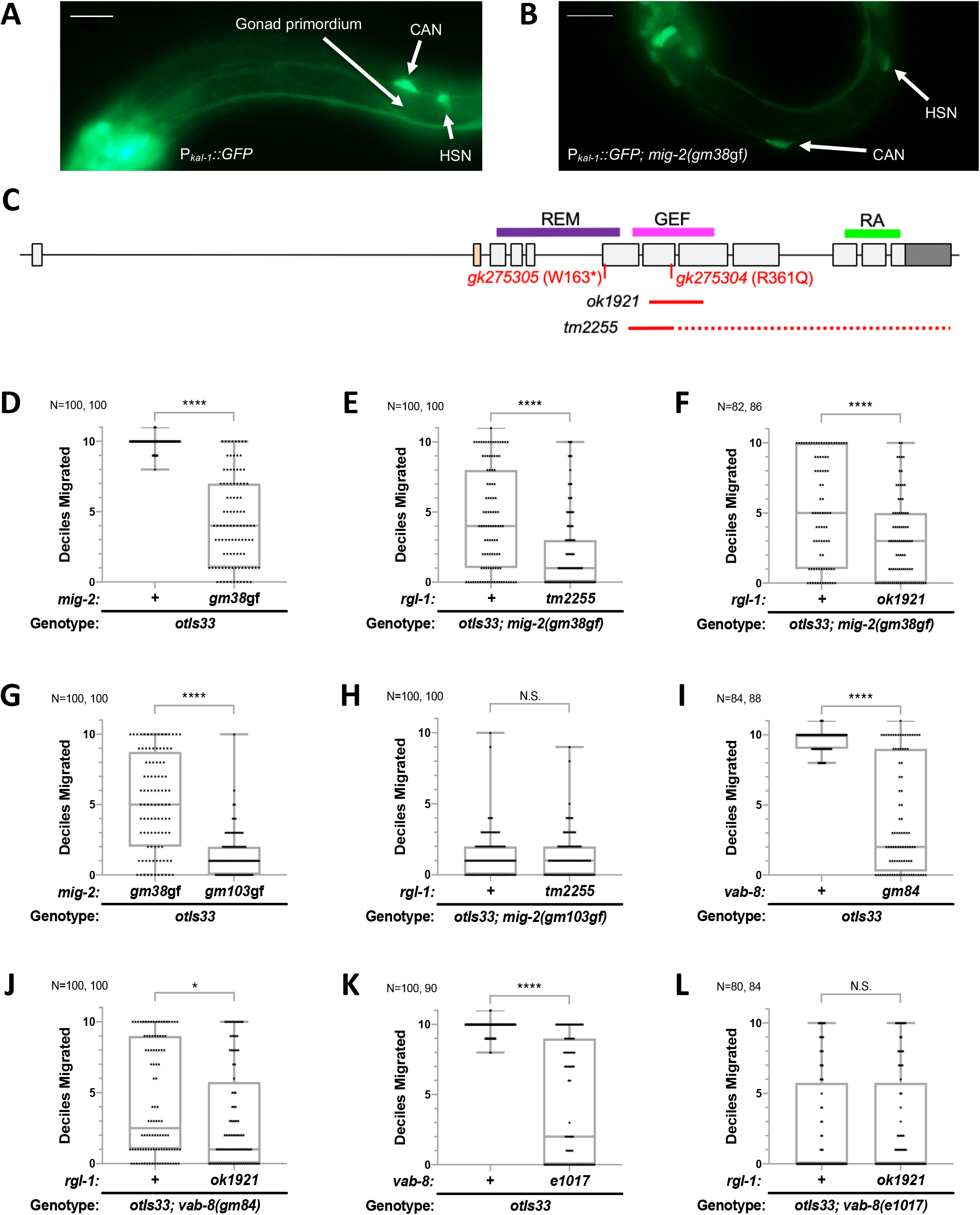
Disruption of RalGEF/RGL-1 enhances mutant defects in CAN migration. **A,B)** Epifluorescent photomicrographs of CAN position at the L1 stage. L1 CANs are positioned just anterior to the gonad primordium, HSNs just posterior. CANs and HSNs, indicated by arrows are bilaterally symmetric, but cells on the other side are out of the plane of focus in these images. Positioning of CANs and HSNs was visualized using the *otIs33* GFP reporter in **A)** wild-type and **B)***mig-2(gm38*gf) backgrounds. Scale bar is 10 μm. **C)** A gene model of *rgl-1* with protein domains indicated above the model and genetic tools indicated below. These include a putative GEF-dead missense allele, *gk275304* (R361Q), nonsense allele *gk275305* (W163*), in-frame deletion *ok1921*, and out-of-frame deletion *tm2255*. The yellow-colored exon encodes 20 residues that are not conserved in other species and is present in ~5% of RNAseq reads. **D-L)** CAN positioning was recorded as decile scores from 0 to 11. 10 indicates the wild-type position, 11 indicates over-migration, and 0 indicates a missing CAN, presumed to have not exited the head but not discernable amongst other GFP-labelled neurons. Any two strains represented by a graph were scored at the same time and were grown on the same OP50 lawn at the same temperature. Positioning of CANs were scored in the *otIs33[*P_*kal-1*_::*GFP]* background unless otherwise noted. **D)***mig-2(gm38*gf) confers a moderate migration defect. **E, F)** The *rgl-1* deletions *tm2255* and *ok1921*, respectively, enhanced the migration defect of *gm38*gf. G) *mig-2(gm103*gf) conferred stronger migration defects than did *mig-2(gm38*gf). **H)** Deletion allele *rgl-1(tm2255)* failed to enhance the defect of *gm103*gf animals. **I, K)** Moderate and strong alleles of *vab-8*, *gm84* and *e1017*, respectively confer CAN migration defects. **J, L)** The rgl-1 deletion allele *ok1921* enhanced the moderate *gm84* but not the strong *e1017* alleles of *vab-8*, respectively. Throughout, each data point is shown. The median is shown by a line, the box indicates 25% and 75% confidence intervals, and bars indicate outlying data. **** represents P<0.0001, *** P<0.001, ** P<0.01, and * P<0.05.

**Figure 2:**
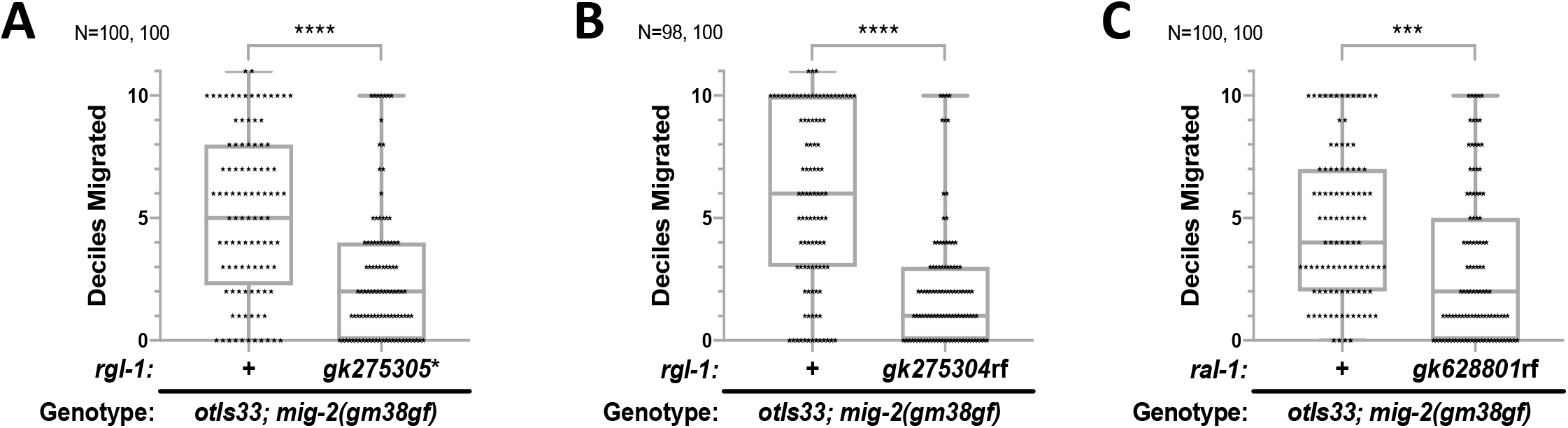
RalGEF signals through Ral to regulate CAN positioning. **A, B)** The *rgl-1* nonsense and GEF-specific alleles, *gk275305\** and *gk275304*rf, respectively, both enhance the CAN positioning defect of *mig-2(gm38*gf). **C)** The *ral-1(gk628801*rf) allele that abrogates RAL-1 signaling also enhances the CAN positioning defect of *mig-2(gm38*gf). The median is shown by a line, the box indicates 25% and 75% confidence intervals, and bars indicate outlying data. **** represents P<0.0001 and *** P<0.001.

**Figure 3:**
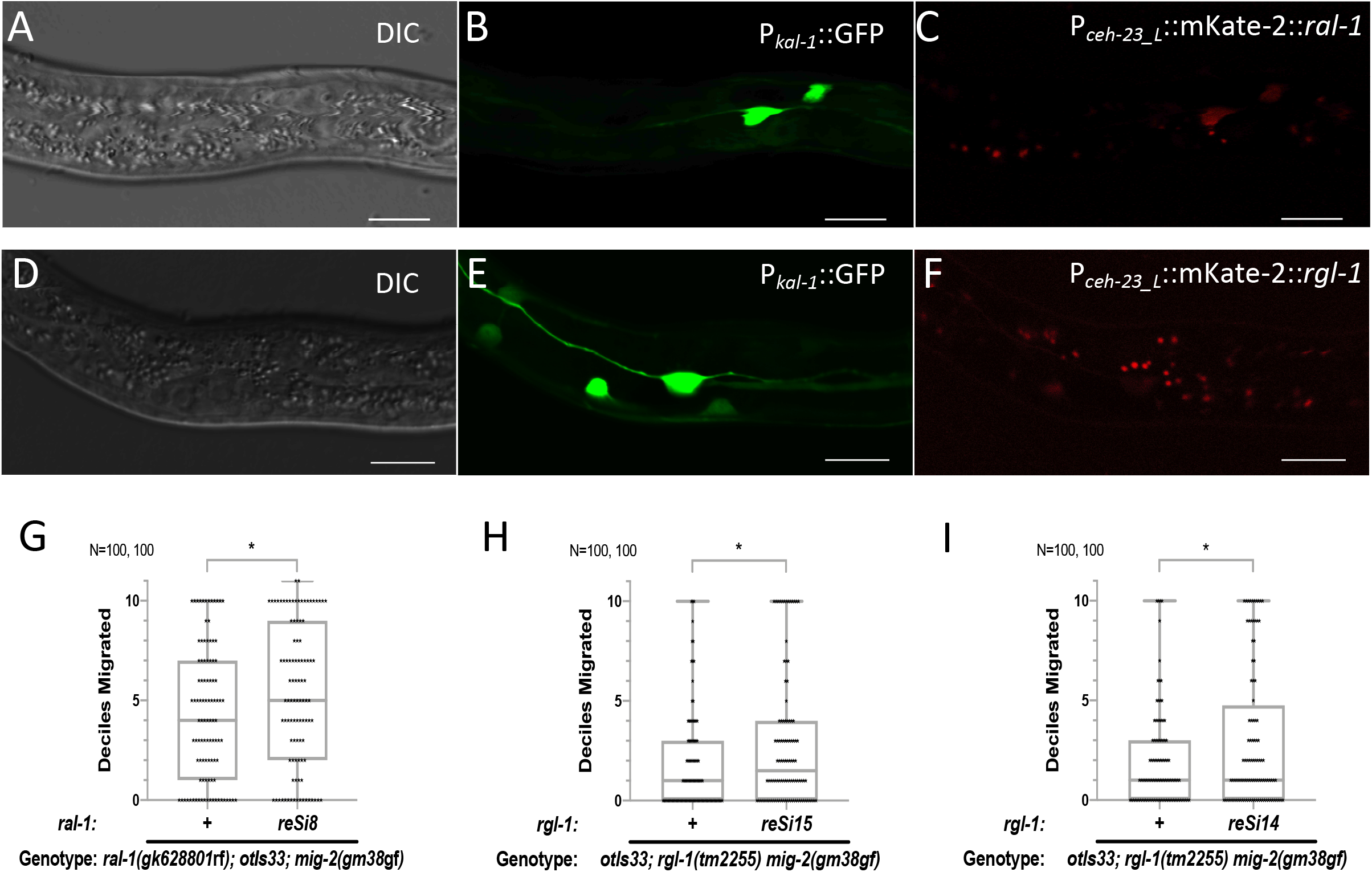
RalGEF>Ral signaling functions cell autonomously to regulate can positioning. **A, B, C)** Photomicrographs of CAN-specific expression of rescuing mKate2::RAL-1 in an animal of genotype *reSi8[P_ceh-23_L_::mK2::ral-1]*; *ral-1(gk628801*rf); *otIs33[P_kal-1_::gfp]* using DIC, green and red channels, respectively. **D,E,F)** Photomicrographs of CAN-specific expression of rescuing mKate2::RGL-1 in an animal of genotype *reSi14[P_ceh-23_L_::mK2::rgl-1]*; *rgl-1(gk275304*rf); *otIs33[P_kal-1_::gfp]* using DIC, green and red channels, respectively. **G)** CAN-specific expression of mK2::RAL-1 partially rescues the enhanced CAN migration phenotype of *gk628801*; *gm38*gf. **H, I)** CAN-specific expression of mK2::RGL-1 partially rescues the enhanced CAN migration phenotype of *gk275304*; *gm38*gf. * represents P<0.05.

### CRISPR/Cas9 genome editing

Insertions for potential transgenic rescue were generated via CRISPR into site I: −5.32., using plasmid as a repair temple. *reSi14[P_ceh-23_L_::mKate2::rgl-1]*, *reSi15[P_ceh-23_L_::mKate2::rgl-1]*, *reSi8[P_ceh-23_L_::mKate2::ral-1]*, and *reSi9[Pceh-23_L::mKate-2::ral-1]*. Gibson Assembly (NEB) was used to generate the CRISPR templates, pJM1 and pJM2 as described above. The presumptive ttTi4348 MosSCI site on Chromosome I was used as the cutting site with crRNA JIM063 used for all four CRISPR alleles.

Alleles *reSi14* and *reSi15* were generated with inner primers JIM059/070 and outer (with homology arms) primers JIM061/071, which were used to amplify pJM1 by PCR. PCR products with long and short homology arms were melted and re-annealed to generate heteroduplexes with long single-stranded overhangs (Dokshin et al., 2018). *S.* pyogenes Cas9 3NLS (250 μg/μL), tracrRNA (100 μg/μL), donor crRNA (28 μg/μL), *dpy-10* crRNA (25 μg/μL), repair template ssODN (10 μM), and dpy-10(cr64) ssODN (600 nM) were microinjected into DV3577 *otIs33[P_kal-1_::GFP]*IV; *rgl-1(tm2255)* animals. Genotyping and sequencing of these *reSi14 and reSI15* CRISPR inserts were performed with JIM064/072/066 (Tm=60°C).

Alleles *reSi8* and *reSi9* were generated with inner primers JIM059/060 and outer primers JIM061/062, which were used to amplify pJM2 by PCR. This was then melted and re-annealed similarly to reSi14 and reSi15 and microinjected into DV2926 *ral-1(gk628801)*III; *otIs33[P_kal-1_::GFP]*IV. Genotyping and sequencing of these *reSi8* and *reSi9* were performed with JIM064/065/066 (Tm=60°C).

### CAN positioning assays

During embryogenesis, the bilaterally symmetrical canal associated neurons (CANs) are born in the head and migrate posteriorly, to a point just anterior to the gonad primordium (Sulston, 1983). In this study, we infer defects in migration of the CANs by their final position upon hatching. To score the position of (CANs), L1 animals were mounted onto slides with a 3% NG agar pad in 5 μL of 2 mg/mL tetramisole/M9 buffer. To track CAN positions, we used the *otIs33[P_kal-1_::gfp]* reporter to observe the final placement of CANs in first larval stage (L1 animals) (Fig. 1A,B). CAN position was analyzed by GFP epifluorescence using a Nikon Eclipse TE2000U microscope. CANs were scored in deciles from zero to eleven. In wild type animals, CANs are positioned immediately anterior to the gonad primordium, designated as the baseline “10” position. In animals with the strongest defects, the CAN was not visible; we would infer them to be located in the cluster of head neurons also labeled by GFP in *otIs33*. Thus, when the CAN on one side was absent, it was scored as a 0. Under-migration was scored based on its location in the L1 animal between the head and gonad primordium, between 1 and 9, and over-migration was scored as an 11. Both left and right CANs were scored in each animal.

### Locomotion assay

Locomotion was assayed as described (Reiner et al., 1999; Reiner et al., 2006). Briefly, young hermaphrodite adults were placed in the center of a 10-cm plate with a three-day old evenly distributed *E. coli* OP50 lawn, and the origin was marked. Animals were allowed to move freely on the plate for 25 minutes at 20°C. The plates were then quickly transferred to −20°C and left for 5 minutes to arrest animal movement. The final location of each animal was marked and the radial distance from the origin to the final point was measured to the nearest half mm. Statistical analysis was performed using Mann-Whitney U-test (see figure legends for P values).

### Confocal microscopy

L1 animals were mounted onto slides with a 3% NG agar pad in 5 μL of 2 mg/mL tetramisole/M9 buffer. Confocal images were captured by A1si Confocal Laser Microscope (Nikon) with 488, 561nm lasers using NIS Elements Advanced Research, Version 4.40 software (Nikon). To ensure consistent exposures the same settings were used in imaging of both CRISPR tagged strains: 488nm (115, 0, 5), 561nm (160, 0, 10) (HV, Offset, Laser Power).

## RESULTS

### Disruption of RalGEF^RGL-1^ enhances mutant defects in CAN positioning

To investigate a possible role of RalGEF>Ral signaling in metastasis, we examined the impact of genetic disruption of RalGEF>Ral on anterior-to-posterior (A>P) cell migrations of GFP-labeled <u>c</u>anal-<u>a</u>ssociated <u>n</u>eurons (CANs; see Methods). The bilaterally symmetrical CANs migrate during embryonic development, after most mitotic cell proliferation has ceased (Forrester and Garriga, 1997; Sulston, 1983). Mutations in RalGEF^RGL-1^ were analyzed in an otherwise wild-type background containing the *otIs33[*P_*kal-1*_::*gfp]* marker for CAN cells (Fig. 1A), or in the *mig-2(gm38*gf) sensitizing mutant background that is partially defective in CAN migrations (Fig. 1B,D; Zipkin et al., 1997)).

Mutations in RalGEF^RGL-1^ are shown in Fig. 1C, based on our previous analysis of RalGEF^RGL-1^ function in patterning of VPC fates (Shin et al., 2018). The out-of-frame deletion *rgl-1(tm2255)* and in-frame deletion *rgl-1(ok1921)* did not confer CAN positioning defects in an otherwise wild-type background (Fig. S1A,B). Both RalGEF^RGL-1^ deletion alleles significantly enhanced the CAN positioning defect of *mig-2(gm38*gf) mutant animals (Fig. 1E,F). Both deletions also enhanced the CAN positioning defect of *mig-2(gm38*gf) as detected using the *lqIs27[*P_*ceh-23*_::*gfp]* reporter, thus controlling for effects of different reporters (Fig. S1H,I). The bilaterally symmetric HSN neurons occupy a mid-body position similar to the CAN neurons, but migrate in the opposite direction, from posterior to anterior (P>A; Desai et al., 1988; Manser and Wood, 1990). *tm2255* also enhanced the defects in positioning of the HSNs, also marked by the *otIs33* reporter (Fig S1J). Thus, RalGEF^RGL-1^ functions in a modulatory role to regulate both A>P and P>A cell migrations.

We additionally examined the effects of deleted RalGEF^RGL-1^ on a second allele of *mig-2*. *mig-2(gm103*gf) confers a significantly stronger positioning defect than does *mig-2(gm38*gf) (Zipkin et al., 1997). *mig-2(gm38*gf) causes an A163V change in RhoG^MIG-2^, a member of the Rho family in the Ras superfamily of small GTPases, that is analogous to the non-canonical A146V weakly activating mutation in Ras (Edkins et al., 2006; Feig and Cooper, 1988; Gripp et al., 2008; Martins-Chaves et al., 2020; Serrano et al., 2016). In contrast, *mig-2(gm103*gf) causes a canonical G16E change, analogous to the maximally activating oncogenic G12D mutation in Ras that prevents inactivation of small GTPases, and is thus of maximum strength (Wennerberg et al., 2005). The phenotype conferred by *gm103*gf is not further enhanced by *rgl-1(tm2255)* (Fig. 1G,H). Thus, either the *rgl-1* interaction with *mig-2* is specific to the non-canonical activating mechanism of A146V change, or cannot further enhance strong defects in CAN position caused by full activation of MIG-2.

To discriminate between these possibilities, we evaluated the impact of RalGEF^RGL-1^ mutation in other mutant backgrounds that confer defective CAN positions. Mutations in KIF26^VAB-8^ (kinesin family member-like) confer defects in CAN migration (Wightman et al., 1996; Wolf et al., 1998). The CAN position phenotype of intermediately defective *vab-8(gm84)* phenotype was enhanced by *rgl-1(ok1921)* (Fig. 1I,J), while that of strongly defective *vab-8(e1017)* was not enhanced by *rgl-1(ok1921)* (Fig. 1K,L). Mutations in EVL^UNC-34^ (<u>E</u>na/<u>V</u>ASP-<u>l</u>ike) also caused moderate defects in CAN positioning (Fleming et al., 2010). The CAN position phenotype of *unc-34(e315)* was enhanced by *rgl-1(ok1921)* and *rgl-1(tm2255)* (Fig. S1C,D,E).

Taken together, our results suggest that enhancement of *mig-2(gm38*gf) phenotypes by mutant RalGEF^RGL-1^ is not specific to *gm38*gf, and is due to the strength of CAN position defect: moderate but not severe defects can be enhanced. Alternatively, these results are consistent with RalGEF^RGL-1^ and Ral^RAL-1^ functioning in the same linear pathway as RhoG^MIG-2^ and KIF26^VAB-8^ and in parallel with EVL^UNC-34^. Alternatively, perhaps multiple signals converge on the same endpoint, defects in which cannot be enhanced.

### RalGEF signaling through Ral regulates migration

In patterning of VPC fates, we previously determined that RalGEF^RGL-1^ performs two opposing functions. Canonical Ras^LET-60^>RalGEF^RGL-1^>Ral^RAL-1^ signaling functions in a modulatory role to promote 2° vulval fate (Shin et al., 2018; Zand et al., 2011), while non-canonical, GEF-independent RalGEF^RGL-1^ functions in a modulatory role to promote 1° vulval fate, possibly as a scaffold for PDK^PDK-1^>Akt^AKT-1^ signaling (Shin et al., 2019). Additionally, Ral^RAL-1^ functions in an essential role in the function of the exocyst complex, which is independent of its role in signaling: complete deletion of Ral^RAL-1^ confers sterility, while signaling defective Ral^RAL-1^ only abrogates signaling activity (Armenti et al., 2014; Shin et al., 2019; Zand et al., 2011). Using reagents we developed to genetically separate functions of RalGEF^RGL-1^, we analyzed whether the role of RalGEF^RGL-1^ in CAN positioning is canonical (GEF-dependent) or non-canonical (GEF-independent).

Both the *rgl-1(gk275305)* W163* nonsense and the *rgl-1(gk275304)* R361Q putative GEF dead mutations (Shin et al., 2019) enhanced the CAN position defect of *mig-2(gm38*gf) (Fig. 2A,B). Neither allele alone conferred defects in CAN position (Fig. S2A,B). The specificity of the *rgl-1(gk275304)* R361Q putative GEF dead mutation is consistent with the function in CAN positioning being GEF-dependent.

The *ral-1(gk628801*rf) R139H mutation alters an Arginine residue that is conserved in all Ras family small GTPases in *C. elegans*, *Drosophila*, and mammals, and abrogated the 2°-promoting function of Ral^RAL-1^ (Shin et al., 2019; Shin et al., 2018). *ral-1(gk628801*rf) does not alter the essential function of Ral, which is likely independent of GEF and GAP function and hence GDP/GTP switching (Shin et al., 2019). *ral-1(gk628801*rf) enhanced the CAN positioning defect of *mig-2(gm38*gf), but conferred no positioning defect as a single mutant (Fig. 2C; S2C). Taken together, these results indicate that it is canonical RalGEF^RGL-1^ activation of Ral^RAL-1^ signaling that contributes to CAN migration.

### RalGEF>Ral signaling functions cell autonomously to contribute to CAN migration

To test whether RalGEF^RGL-1^>Ral^RAL-1^ signaling functions cell autonomously to modulate CAN migrations, we performed experiments to test transgenic rescue of mutant enhancement of *mig-2(gm38*gf) CAN migration defects. We used CRISPR to insert sequences containing P_*ceh-23_L*_::*mKate2*::*ral*-1 into a safe harbor position on Chromosome I (map position −5.32; see Methods). We confirmed by red fluorescence that transgenic mKate::RAL-1 was expressed in CANs, labelled in *otIs33* by coincident *P_kal-1_::gfp* (Fig. 3A-C). This construct partially rescues the enhancement of CAN migration defects by *ral-1(gk628801*rf) in a *mig-2(gm38*gf) background (Fig. 3G). We performed similar rescue analysis with single copy wild-type mKate2::RGL-1 in the *rgl-1(tm2255) mig-2(gm38*gf) background (see Methods). We confirmed by red fluorescence that transgenic mKate::RGL-1 was expressed in CANs, labelled in *otIs33* by coincident *P_kal-1_::gfp* (Fig. 3D-F). P_*ceh-23_L*_::*mKate2*::*rgl*-1 partially rescued the enhancement of *gm38*gf by mutant *rgl-1* (Fig. 3H,I). These results are consistent with RalGEF^RGL-1^ and Ral^RAL-1^ functioning cell autonomously in the CANs to contribute to their final positioning. But partial rescue from single copy inserts is also consistent with more complicated interpretations, possibly including non-autonomy.

### Ral^RAL-1^ is expressed in the CANs

We previously tagged endogenous Ral^RAL-1^ and observed apparently ubiquitous expression, to the degree we could assess that. To specifically determine whether Ral^RAL-1^ is expressed CANs, we co-imaged *ral-1(re218[mKate2::RAL-1])* with the *otIs33[Pkal-1::gfp]* marker of CANs, using confocal microscopy (Fig 4A-C). Because the CANs are positioned away from other cells, we were able to visualize RAL-1 expression in the CAN during larval development.

**Figure 4:**
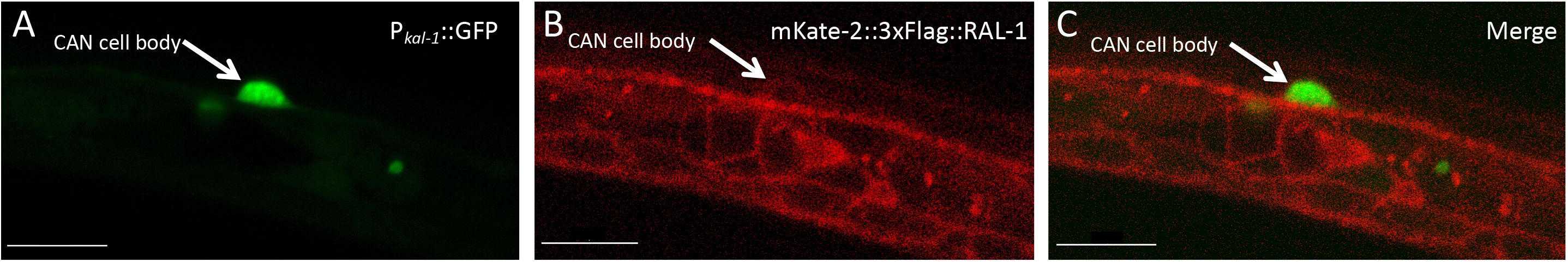
Ral^RAL-1^ is expressed in the CANs. Confocal micrographs. **A)** CAN labeled in green by *otIs33[P_kal-1_::gfp]*. **B)** red-tagged RAL-1, *ral-1(re218[mKate2::2xHA::RAL-1])*. **C)** Merged images of *ral-1(re218[mKate2::RAL-1]); otIs33[Pkal-1::gfp]*. Arrows indicates the CAN. Scale bars = 10 μm.

### Disruption of RalGEF enhances defects in general nervous system function and development

We observed that *rgl-1(ok1921) mig-2(gm38*gf) and *rgl-1(tm2255) mig-2(gm38*gf) double mutant animals had significantly more severe locomotion defects compared to the *mig-2(gm38*gf) single mutant. Because that CAN neurons are not known to function in regulating motility, this observation suggests that other aspects of nervous system anatomy or function require RalGEF^RGL-1^ function. We quantified enhancement of the locomotion defect conferred by *gm38*gf using a circumferential locomotion assay and found that deletion of RalGEF^RGL-1^ increased severity of locomotion defects conferred by *mig-2(gm38*gf) (Fig. 5A; see Methods).

**Figure 5:**
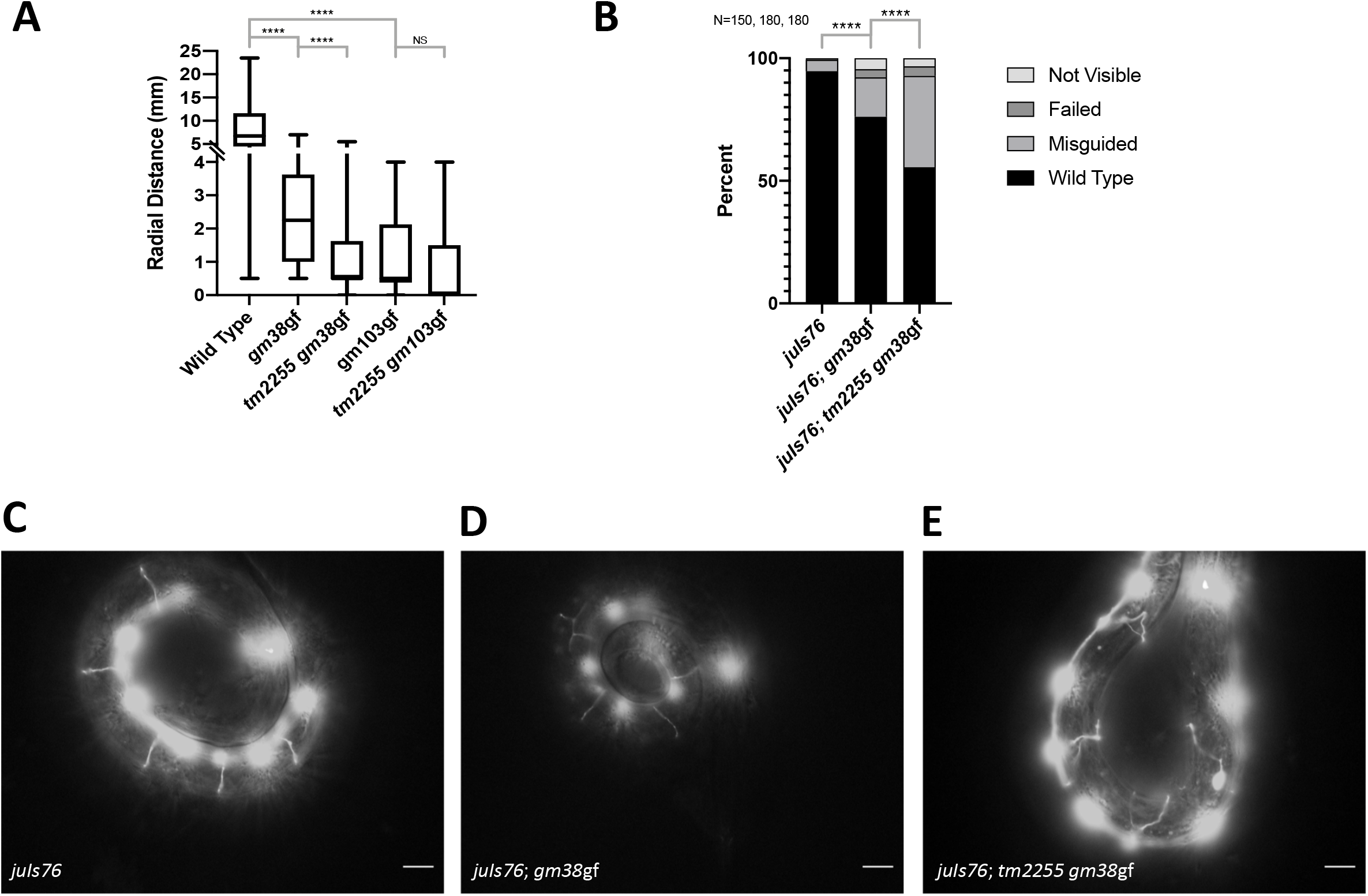
Deletion of RalGEF enhances defects in general nervous system function and development. **A)** Measurement of radial locomotion (see Methods) shows that genetic interactions controlling locomotion reflect those observed for CAN positioning in Figure 1. *mig-2(gm103*gf) is more severely defective in locomotion than is *mig-2(gm38*gf), and that *rgl-1(tm2255)* enhances *gm38* but not *gm103.* **B)** A marker of DD and VD axons, *juIs76[*P_*unc-25*_::*GFP*], revealed that dorsoventral axon guidance was defective in *mig-2(gm38*gf) animals and was enhanced by *rgl-1(tm2255)*. **C, D, E)** Epifluorescence photomicrographs of circumferential axon migration in wild-type, *gm38* and *tm2255 gm38* animals, respectively. Scale bar = 10 μm. **** represents P<0.0001.

The DD and VD motor neurons are key regulators that coordinate locomotion. In order to evaluate the neuronal anatomy of these motor neurons, we visualized the circumferential, dorsal-ventral (D-V) axons of DD and VD motor neurons in the wild type, single and double mutants using the *juIs76[P_unc-25_::GFP]* reporter (Fig. 5B-E; see Methods; (Jin et al., 1999)). *mig-2(gm38*gf) conferred inappropriate deviations of axon pathfinding from the wild type dorsal-ventral axis (Fig. 5B-D). This defect was significantly enhanced by *rgl-1(tm2255)* (Fig. 5B,D,E). These results suggest that RalGEF^RGL-1^ regulates growth cone guidance as well as cell migration, that RalGEF^RGL-1^ functions in a modulatory capacity to regulate guidance events along both A-P and D-V axes of the animal, and perhaps that RalGEF^RGL-1^ functions generally to sculpt the nervous system in collaboration with other signals.

### Ral^RAL-1^ functions as a permissive cue in CAN migration

RalGEF^RGL-1^>Ral^RAL-1^ is necessary for maximal migration of CAN cells when other guidance signals are perturbed. The contribution of RalGEF^RGL-1^>Ral^RAL-1^ signaling to CAN migration could comprise either a permissive or instructive cue. If Ral^RAL-1^ functions as a permissive signal, a single outcome is evoked by Ral^RAL-1^ signal, in this case an incremental advance in CAN migration. Thus, we would predict that constitutively activated Ral^RAL-1^ would not confer migration defects in CANs, because sufficient signal is transduced by wild-type Ral^RAL-1^ to satisfy a functional requirement. In contrast, if Ral^RAL-1^ functions as an instructive signal, Ral would function continually to provide positional information to the cell. Excess Ral^RAL-1^ activation would be expected to perturb CAN positioning as much or more than insufficient Ral^RAL-1^ signaling, and hence constitutively activated Ral^RAL-1^ would be predicted to enhance the CAN positioning defect conferred by *mig-2(gm38*gf). An example of an instructive cue in CAN migration is RhoG^MIG-2^ itself: loss of *mig-2* function causes no discernable defects in CAN positioning, while the moderate and strong gain-of-function mutations dramatically perturb CAN position (Zipkin et al., 1997). Another example of an instructive cue in CAN migration are the FGF^EGL-17^ growth factor and the FGFR^EGL-15^ receptor. Constitutively activated FGFR^EGL-15^ in the CANs enhances defective CAN positioning in a sensitized background, as does heat shock of its ligand, FGF^EGL-17^, during the period of migration (Fleming et al., 2005).

Using CRISPR, we previously generated *ral-1(re160*gf*[mKate2^2xHA::ral-1(G26V)])* (analogous to the maximally activated G12V in Ras numbering), which is both constitutively activated and tagged at the N-terminus with a mKate2 fluorescent protein (Shin et al., 2018). We also generated *ral-1(re218[mKate2^2xHA::ral-1])*, which is tagged but otherwise wild type. We found that *ral-1(re160*gf) relative to *ral-1(re218)* did not alter CAN position in the *mig-2(gm38*gf) or wild-type background (Fig. 6A; S3A,B). These observations suggest that RalGEF^RGL-1^>Ral^RAL-1^ functions as a permissive signal, with a simple requirement for function that is neither spatial nor dose-sensitive.

**Figure 6:**
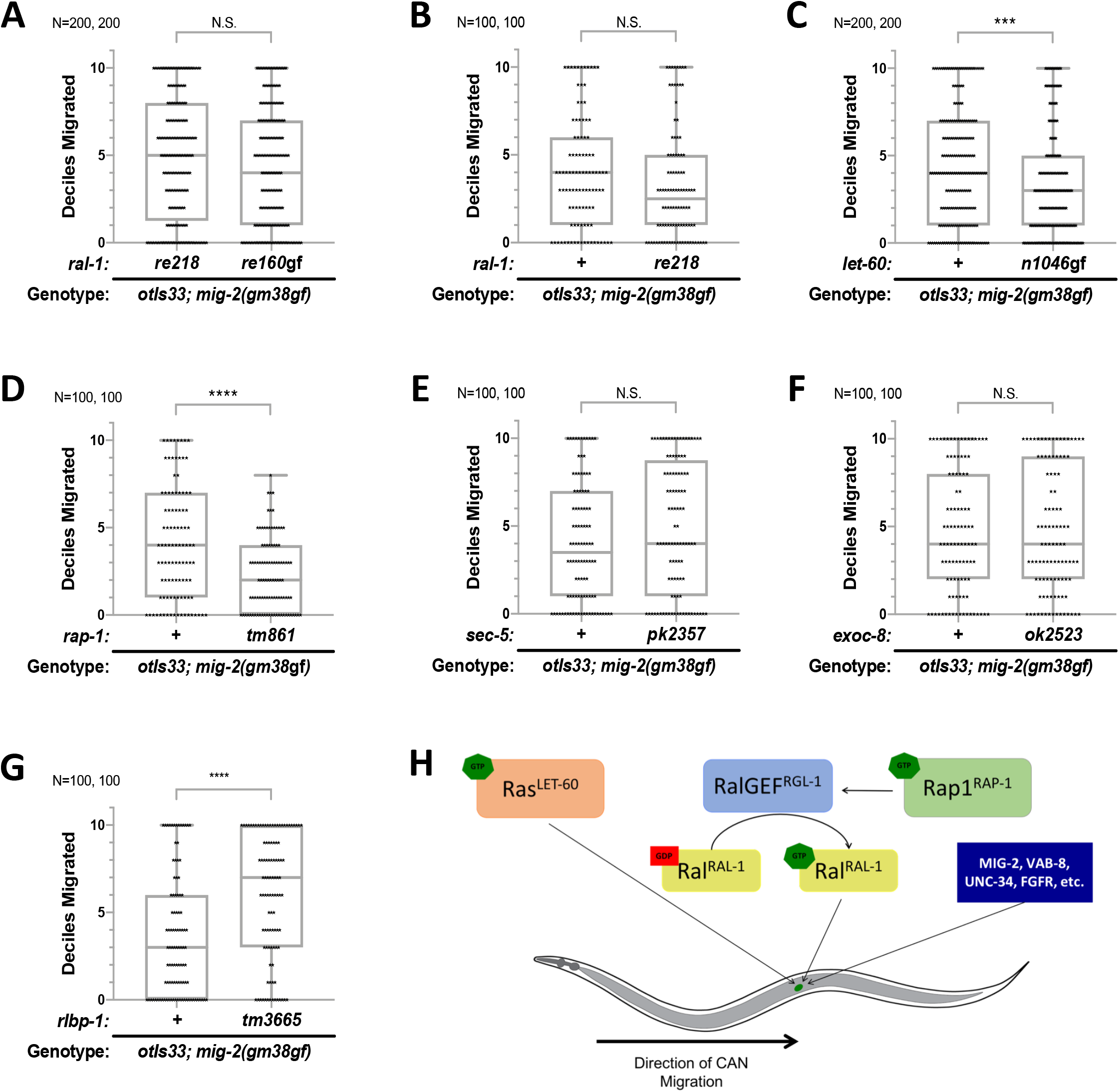
Ral^RAL-1^ functions as a permissive cue while Ras^LET-60^ functions as an instructive cue in CAN migration. **A,B)** Constitutively active *ral-1(re160gf) and w*ild-type *ral-1(re218)*, both tagged with fluorescent protein and epitope, fail to enhance the CAN positioning defect conferred by *mig-2(gm38*gf*). let-60(n1046*gf), which causes constitutive activation, enhances the CAN positioning defect of *mig-2(gm38gf)*. **D)** Deletion of rap-1, *rap-1(tm861)* enhanced the positioning defect of *gm38*gf. **E)** Reduction of function of putative Ral effector *sec-5* function did not alter CAN positioning in the *gm38*gf background. Non-Green homozygous *sec-5(pk2357*rf) animals from heterozygous mothers in which the *sec-5* mutation was balanced by GFP-labelled chromosomal inversion, *mIn1*. **F)** Deletion of putative Ral effector Exo84, *exoc-8(ok2523)*, did not alter CAN positioning in the *gm38*gf background. G) Deletion of putative Ral effector RalBP1 by *rlbp-1(tm3665)* unexpectedly suppressed the migration defect of *gm38*gf. The median is shown by a line, the box indicates 25% and 75% confidence intervals, and bars indicate outlying data. **** represents P<0.0001, *** P<0.001, ** P<0.01, and * P<0.05. **H)** A model for the function of parallel Rap1^RAP-1^>RalGEF^RGL-1^>Ral^RAL-1^ and Ras^LET-60^ signals controlling migration of the CANs, as well as other parallel signals.

### Ras^LET-60^ functions as an instructive cue in CAN migration

Since the Ras>RalGEF>Ral signaling module is found in various systems, including *C. elegans* patterning of vulval precursor cell (VPC) fates, we expected that Ras^LET-60^, like Ral^RAL-1^, would function as a permissive cue. We used the *let-60(n1046*gf) mutation, which causes an activating G13E mutation in Ras^LET-60^ sufficient to transform presumptive 3° vulval cells into 1° cells (Beitel et al., 1990; Han et al., 1990), to test the contribution of Ras^LET-60^ to CAN migration. *let-60(n1046*gf) conferred an enhanced defect in CAN positioning in the *mig-2(gm38*gf) background (Fig. S3D). Since the P value of 0.033 was of borderline significance, we were concerned about the possibility of type I error (a false positive). Consequently, we repeated scoring of *gm38*gf vs. *n1046*gf; *gm38*gf animals with twice as many animals scored (N = 200 each rather than N=100), which validated the significance of the result (Fig. 6C). *let-60(n1046*gf) alone did not alter CAN position (Fig. S3C). This result is striking because *let-60(n1046*gf) G13E is predicted to be only moderately activating. In contrast, the *ral-1(re160*gf) G26V (G12V in Ras numbering) is predicted to be maximally activating. (Both G12V and G13D oncogenic mutations are present in human H-, N- and K-Ras (Pylayeva-Gupta et al., 2011; Reiner and Lundquist, 2018)). Thus, if anything we would expect *re160* to confer a stronger defect than *n1046*.

We propose three possible models to explain these observations. First, Ras^LET-60^ is instructive and signals through an effector other than RalGEF^RGL-1^>Ral^RAL-1^. This model would perhaps necessitate that RalGEF^RGL-1^>Ral^RAL-1^ is activated by a Ras-like protein other than Ras^LET-60^, possibly Rap1^RAP-1^ (Rasmussen et al., 2018); see below), and that this protein functions permissively, not instructively, to activate RalGEF^RGL-1^>Ral^RAL-1^. Second, Ras^LET-60^ signals through two effectors, like both RalGEF^RGL-1^>Ral^RAL-1^ and Raf^LIN-45^, and that both of them together reveal an instructive function where one does not. Third, Ras^LET-60^ could mediate two signals, one instructive and one permissive. We cannot test the latter two models, but we can test the first model.

### Rap1^RAP-1^ rather than Ras^LET-60^ activates the RalGEF^RGL-1^>Ral^RAL-1^ signal

The effects of *let-60(n1046*gf) enhances the migration defect of *mig-2(gm38*gf) while *ral-1(re160*gf) does not (see Fig. 6A vs. Fig. 6C). One possible interpretation is that a small GTPase other than Ras^LET-60^ signals through RalGEF^RGL-1^>Ral^RAL-1^ to control migration of CANs. The Ras and Rap1 (<u>Ra</u>s <u>p</u>roximal) small GTPases share identical effector binding sequences (Rasmussen et al., 2018; Reiner and Lundquist, 2018). In *Drosophila*, Rap1 has been implicated in binding and activating RalGEF (Carmena et al., 2011; Mirey et al., 2003). By yeast two hybrid screen, *C. elegans* RAP-1 was found to bind to RalGEF^RGL-1^ (Frische et al., 2007). Consequently, we used genetics and our *mig-2(gm38*gf) CAN migration assay to test whether *C. elegans* Rap1^RAP-1^ is likely to function upstream of RalGEF^RGL-1^>Ral^RAL-1^.

The *rap-1(tm861)* deletion is a null mutation in Rap1^RAP-1^ (Rasmussen et al., 2018). We observed that *rap-1(tm861)* enhances the CAN migration defect of *mig-2(gm38*gf) (Fig. 6D). This result is consistent with a Rap1^RAP-1^>RalGEF^RGL-1^>Ral^RAL-1^ signal functioning in parallel with Ras^LET-60^ to modulate positioning of CANs.

### No evidence for canonical oncogenic Ral effectors functioning downstream of Ral in positioning of CANs

The three oncogenic effectors of mammalian Ral are RalBP1, Sec5 and Exo84 (Gentry et al., 2014; Kashatus, 2013). Each of these is conserved in a single *C. elegans* ortholog (Frische et al., 2007; Zand et al., 2011). We would expect mutant effectors to phenocopy mutant Ral if they are downstream of Ral in the CAN positioning pathway.

The *ok2328* deletion in Exo84^EXOC-8^ and the *pk2357* late stop allele in Sec5^SEC-5^ failed to confer defects in CAN positioning in an otherwise wild-type background (Fig. S3F,G). either *ok2328* nor *pk2357* enhanced the CAN positioning defects of *mig-2(gm38*gf) (Fig. 6E,F). Thus, the role of canonical Ral effectors remains unclear in control of CAN positioning by Ral^RAL-1^.

We tested the ability of *tm3665* deletion in RalBP1^RLBP-1^ to enhance the CAN positioning defect of *mig-2(gm38*gf). Unexpectedly, deletion of RalBP1^RLBP-1^ suppressed, rather than enhanced, the positioning defect of *gm38*gf, but had no effect in a wild-type background (Fig. 6G; S3H). Mammalian RalBP1 functions as a GAP to inhibit Cdc42 and Rac, both regulators of cytoskeletal dynamics. Among many possible explanations of our observation, perhaps the function of RalBP1 is subject to complex spatial regulation by Ral, Ral uses additional effectors, or multiple functions of RalBP1^RLBP-1^ contribute to the observed genetic interaction (Neel et al., 2012). Taken together, these results leave unresolved the role of canonical Ral effectors in positioning of the CANs.

## DISCUSSION

We have observed that the RalGEF^RGL-1^>Ral^RAL-1^ signaling module, originally defined as an oncogenic effector downstream of Ras, broadly regulates morphogenesis of the nervous system. This function appears to be strictly modulatory: we did not detect defects in an otherwise wild-type background upon disruption of RalGEF^RGL-1^>Ral^RAL-1^ function. Instead, we detected enhancement of defects only when other components necessary to cell migration and nervous system development have already been mutated. Our main assay is the positioning of the CANs after embryonic posterior-to-anterior migration, but we also analyze positioning of the HSNs after embryonic anterior-to-posterior migration, general locomotion of the animal, and growth cone guidance of circumferential axon commissures (dorsal-ventral axis). Based on these results, we propose that the RalGEF^RGL-1^>Ral^RAL-1^ signal functions broadly in the animal to modulate neuronal and axonal movements. We additionally found that Ras^LET-60^ and Rap1^RAP-1^ likely govern distinct signaling cascades, with Rap1^RAP-1^ more consistent as the upstream activator of RalGEF^RGL-1^>Ral^RAL-1^ signaling. This study provides an advance in our understanding of both the development of nervous system and the functions of signal transduction of Ras family small GTPases and their effectors.

### The role of Ras and its sibling small GTPases

Unexpectedly, we observed that the function of Ras^LET-60^ is non-equivalent to that of Ral^RAL-1^: a putatively moderate gain-of-function mutation in Ras^LET-60^ confers enhancement of migration defects while a putatively strong gain-of-function mutation in Ral^RAL-1^ does not. A caveat to this result is that Ras^LET-60^, but not Rap1^RAP-1^ and RALGEF^RGL-1^, is essential for development in *C. elegans* (with the *gk628801* R139H mutation we were also able to determine that the signaling activity of Ral^RAL-1^ is not essential for viability). We were unable to evaluate loss-of-function of Ras^LET-60^ because we could not construct double mutant combinations between strong reduction-of-function or dominant negative mutations in Ras^LET-60^ and mutations in genes regulating cell migration. Yet the differing roles of constitutively activated Ras^LET-60^ vs. Ral^RAL-1^ suggest they are functionally distinct. However, we cannot rule out the possibility that Ras coordinates both Raf and RalGEF>Ral signals in this process.

The primary association of RalGEF>Ral signaling in the literature is with Ras (Gentry et al., 2014; Kashatus, 2013), including by our group in the modulatory 2°-promoting signal in support of Notch^LIN-12^ during developmental patterning of the vulval precursor cells, where Ras dynamically switches effectors from Raf to RalGEF>Ral (Shin et al., 2018; Zand et al., 2011). In addition to Rap1A and Rap1B subfamily in mammals, two other subfamilies also contain effector loops identical to that of proto-oncogenic mammalian Ras proteins (H-Ras, N-Ras, K-Ras4A and K-Ras4B): R-Ras (R-Ras1 and R-Ras2) and M-Ras. Thus, based on sequence identity in this key effector binding loop, nine functionally distinct proteins in mammals have the potential to interact in some way with overlapping sets of effectors. The question of whether this happens *in vivo* or how it is regulated has not been investigated via perturbation of endogenous genes. The *C. elegans* and *Drosophila* genomes each encodes four of these proteins: Ras^LET-60^, Rap1^RAP-1^, R-Ras^RAS-1^ and M-Ras^RAS-2^ in *C. elegans* parlance. Of these, all in humans, *Drosophila* and *C. elegans* have identical effector-binding loops with the exception of *C. elegans* M-Ras^RAS-2^, which harbors a conservative change in the core effector binding loop (Reiner and Lundquist, 2018; Wennerberg et al., 2005).

Our finding that deletion of Rap1^RAP-1^ phenocopies mutations in RalGEF^RGL-1^ and Ral^RAL-1^ provides a novel role for Rap1^RAP-1^ in development. This model is supported by instances in *Drosophila* of a Rap1>RalGEF>Ral signal (Carmena et al., 2011; Mirey et al., 2003). Consequently, Rap1 represents a compelling alternative to Ras^LET-60^ as the immediately upstream trigger of RalGEF^RGL-1^>Ral^RAL-1^ signaling.

This phenomenon validates the possibility of interchangeability of close-Ras relatives in different developmental contexts. What advantage would such a regulatory switch confer? First, different GTPases of ostensibly identical biochemical functions can confer distinct subcellular localization due to differences in the C-terminal hypervariable region and prenylated C-terminal tail. These sequences are known to be important for function and confer localization to different subcellular compartments, but the rules governing distinctions between them are not well understood (Kotti, 2018). But GTPase activation in different subcellular compartments could confer different activities during development, as they encounter different effectors to different results in different subcellular locales. Also, upstream positive regulation by different GEFs and negative regulation by different GAPs could mediate different input from diverse receptor types. Some overlap but also some independence is described with GEFs and GAPs for Ras and its closest relative, Rap1 (reviewed in (Raaijmakers and Bos, 2009)). And how is engagement with effectors regulated? A scaffold Ras-Raf interaction, SOC-2/SUR-8, was originally discovered in *C. elegans* and has been shown to be sufficient to induce RASopathy spectrum birth defects when mutated in humans, as are members of the Ras signaling network (Bustelo et al., 2018; Cordeddu et al., 2009; Selfors et al., 1998; Sieburth et al., 1998). Such a molecule could theoretically orchestrate specificity of Ras group interactions with effectors. But scaffolds have not been described for other players named above, and so this mechanism remains speculative. Our study may establish a platform for *in vivo* analysis of such questions.

### The complexity of signals regulating migrations and other morphogenetic events

Using the *vab-8(gm84)* mutant background, another group found no impact of either *let-60(n1046*gf) or a putative null mutation segregating from a heterozygous mother (Tai et al., 2005). (Notably, mutation of the Grb2^SEM-5^ SH2-SH3 adaptor protein did enhance the CAN migration defect in *vab-8(gm84)* animals, illustrating that such adaptors do more than just signal through Ras (Belov and Mohammadi, 2012)). The failure of *let-60(n1046*gf) to enhance *vab-8(gm84)* phenotypes underscores the potentially byzantine network of signals that control cell migration and other migratory events like guidance of growth cones during neurite outgrowth. We hypothesize that genetic backgrounds are a critical factor in revealing functions. For example, in *C. elegans* KIF26^VAB-8^ confers defects only in posterior-directed cell migrations like the CANs by regulation of the Robo receptor system (Watari-Goshima et al., 2007; Wightman et al., 1996; Wolf et al., 1998). Accordingly, we would not expect deficits in RalGEF>Ral signaling to intersect with mutant KIF26^VAB-8^ in other than posterior-directed events. Thus, mutant KIF26^VAB-8^ may not be the ideal background in which to detect effects outside of the CANs by perturbation of RalGEF>Ral signaling, while *mig-2(gm38*gf) was fortuitously sensitive to such perturbations. Consequently, extensive genetic analysis to weave these separate signaling axes into a coherent whole is well beyond the scope of the present study. And thus Rap1^RAP-1^>RalGEF^RGL-1^>Ral^RAL-1^ remains an orphan signal transduction module without known associated receptor function.

Many other signals have been defined in this system. The Rho^RHO-1^, Rac^CED-10^, and Cd42^CDC-42^ Rho family small GTPases, known to control cytoskeletal dynamics as the “business end” of signal transduction during cell migration and neurite outgrowth (Hall, 2005), function throughout these processes during sculpting of the *C. elegans* nervous system, *e.g.* (Alan et al., 2018; Demarco et al., 2012; Gujar et al., 2019) (reviewed in (Reiner and Lundquist, 2018)). Seminal work established the netrin system as playing a central role in D-V oriented events (Hedgecock et al., 1990). FGFR^EGL-15^ has also been shown to function as an instructive cue in CAN migration (Tai et al., 2005). The Ror^CAM-1^ non-canonical receptor tyrosine kinase plays a central role in CAN migrations, perhaps through interactions with Wnt (Forrester et al., 1999; Forrester et al., 2004; Wang and Ding, 2018). Wnt signaling has been shown to function in A-P cell migrations like the CANs as well as A-P polarity of vulval 2° cell lineages (Green et al., 2008). EVL^UNC-34^ functions with the Wnt signaling system to control such A-P events regulated by Wnts (Fleming et al., 2010). Yet even such genetic approaches are limited. For example, double mutants between EVL^UNC-34^ and WASP^WVE-1^ are synthetically lethal due to failure of embryonic gastrulation and morphogenesis (Withee et al., 2004). Yet optimal use of epistasis analysis in generally agreed to require null mutants (Avery and Wasserman, 1992), which is frequently not feasible. A view of this field with one cell type, the migrating Q neuroblasts and descendants, has been reviewed (Rella et al., 2016). But a comprehensive view remains elusive because of the complexity of such spatial morphogenetic events and the extensive redundancy among signaling systems. We are even unsure of whether relationships among different signaling modules retain the same relationships between different developmental events, or whether they are “mix and match” like so much of signaling in developmental biology.

## CONCLUSIONS

In conclusion, we found a role for the exclusive signaling dyad, RalGEF>Ral, in different migratory processes in the nervous system. Wherever analyzed, this signal appears modulatory rather than central to developmental processes; loss of function mutant animals appear wild type. This signal is permissive in CAN migration, in contrast to the instructive Ras^LET-60^ signal. We thus were led to identify a role for the close relative of Ras, Rap1^RAP-1^, in cell migration. We hypothesize that Rap1^RAP-1^ functions to activate RalGEF>Ral in the context of CAN migration, while Ras^LET-60^ constitutes a parallel signal. This study introduces new characters to the cast controlling architecture of the nervous system, finds a role for RalGEF>Ral signaling in morphogenetic events, and also establishes developmental model for RalGEF>Ral in invasion and metastasis using a simple invertebrate model experimental system.

## ACKNOWLEDGEMENTS

This work was supported by NIH grant R01GM121625 and John Templeton Foundation Grant ID# 61099 to D.J.R. Some strains were provided by the CGC, which is funded by NIH Office of Research Infrastructure Programs (P40 OD010440). Wormbase was used regularly.

---

**Figure S1:**
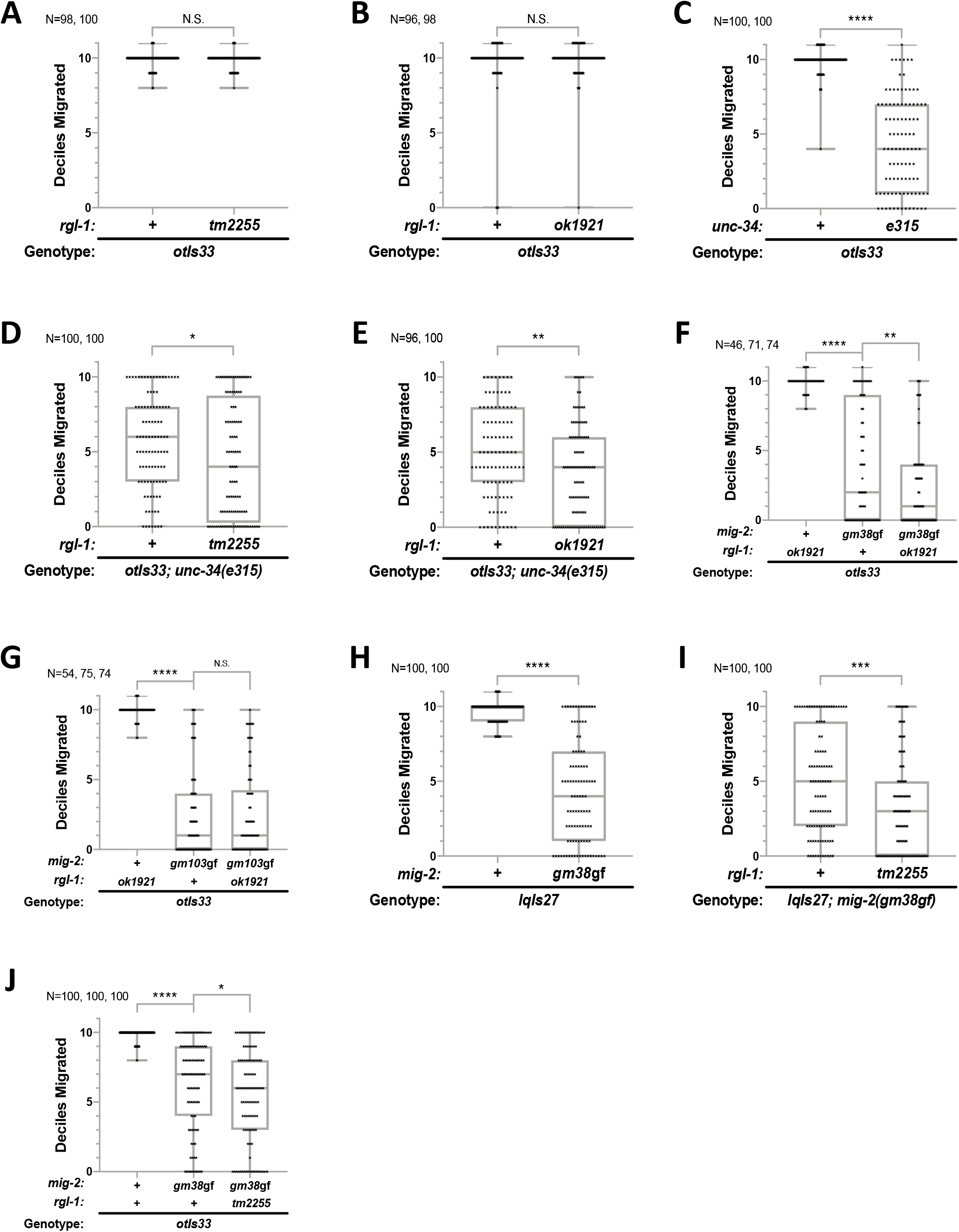
Deletions in *rgl-1* do not cause significant defects in CAN migration as single mutants, and an additional marker and background for CAN migration. **A, B)** Single mutant *tm2255* and *ok1921*, respectively, did not confer defects in CAN positioning. Rare CANs were scored as being mis-positioned in the head. But in these backgrounds with no migration defects, it is likely that the cells were missing, perhaps through mis-specification. It is unknown whether this very low penetrance defect is associated with *rgl-1* or other lesions in these genetic backgrounds. **C, D, E)** The *unc-34(e315*) mutation conferred a defect in CAN positioning that was enhanced by *rgl-1(tm2255*) (D) and *ok1921*. **F, G)** Preliminary results performed performed by D.J.R over a decade before submission reflect that same genetic interactions that were found later. **H, I)** A distinct marker of CAN positioning, *lqIs27[*P_*ceh-23*_::*GFP*], revealed the same effects of *mig-2(mg38*gf) and *rgl-1(tm2255)* as did *otIs33*. **J)***mig-2(gm38*gf) and *rgl-1(tm2255)* impact HSN positioning similarly to how they impact CAN positioning, also marked by *otIs33*. Throughout, each data point is shown. The median is shown by a line, the box indicates 25% and 75% confidence intervals, and bars indicate outlying data. **** represents P<0.0001, *** P<0.001, ** P<0.01, and * P<0.05.

**Figure S2:**
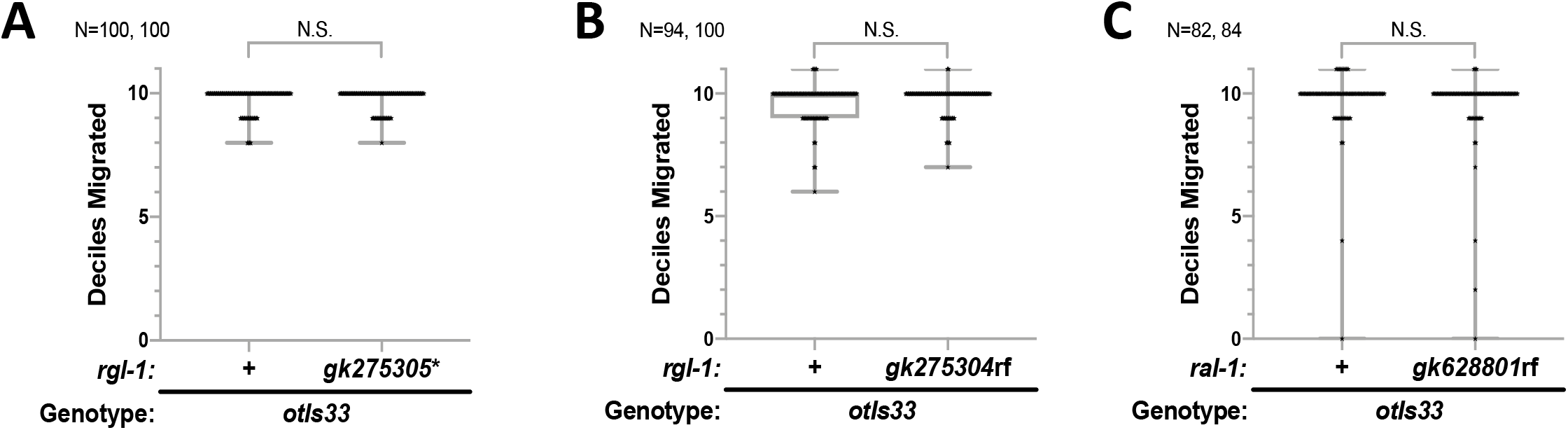
Perturbation of RalGEF->Ral signaling alone does not confer CAN migration defects. **A, B, C)***rgl-1(gk275305*)*, *rgl-1(gk275304*rf), and *ral-1(gk628801*rf) as single mutants fail to confer defects in CAN positioning.

**Figure S3:**
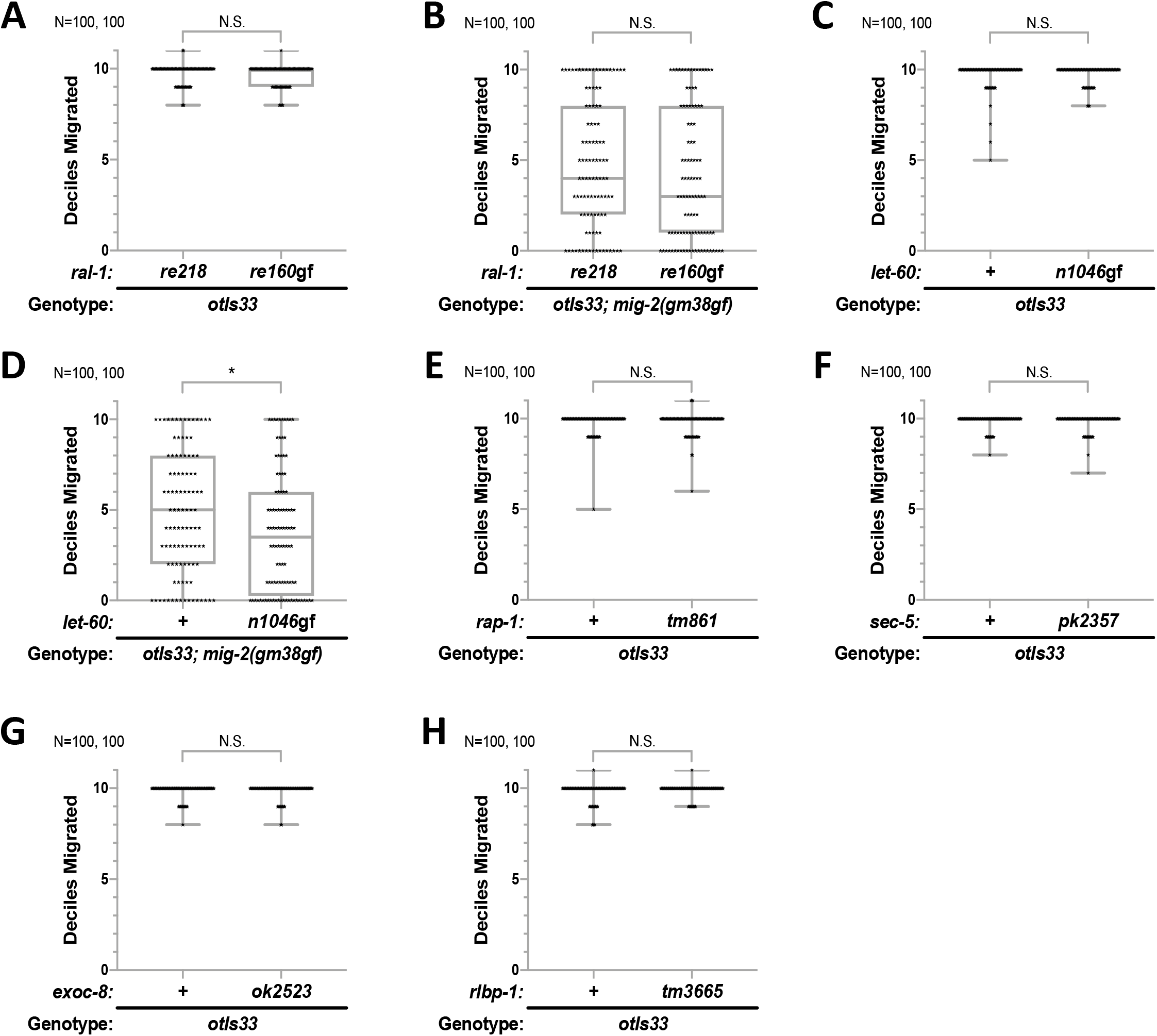
Single mutants for *ral-1* and *let-60* do not confer CAN positioning defects. **A)***ral-1* alleles *re218* and *re160*gf do not confer CAN positioning defects as single mutants. **B)** Original scoring of *re218* vs. *re160gf* in a *mig-2(gm38*gf) background (N =100 each rather than the 200 each presented in Fig. 4). The difference was not significant, but the p-value of 0.144 suggested the possibility of a Type II error (false negative), so we re-tested with a large target N and found a significant difference (presented in Fig. 4). **C)***let-60(n1046*gf) does not confer CAN positioning defects as a single mutant. D) Original data of *let-60(n1046*gf) in a mig-2(gm38gf) mutant background. With a p-value of 0.033, were concerned about a Type I error (false positive), and so we re-tested with a larger target N and found the result to be non-reproducible (presented in Fig. 4C). **E-H)** Single mutants for *rap-1(tm861*)*, sec-5(pk2357*rf), *exoc-8(ok2523)*, and *rlbp-1(tm3665)*, respectively do not confer a significant defect in CAN positioning. Non-Green homozygous *sec-5(pk2357*rf) animals from heterozygous mothers in which the *sec-5* mutation was balanced by GFP-labelled chromosomal inversion, *mIn1*. The median is shown by a line, the box indicates 25% and 75% confidence intervals, and bars indicate outlying data. **** represents P<0.0001, *** P<0.001, ** P<0.01, and * P<0.05.

**Supplementary Table 1:** Strains.

| Strain<sup>a</sup> | Genotype | Application |
| --- | --- | --- |
| CB1017 | *vab-8(e1017)* V | Starting reagent |
| DV2144 | *otIs33[P<sub>kal-1</sub>::GFP]* IV; *mig-2(gm103gf)* X | Fig. 1G |
| DV2148.2 | *otIs33[P<sub>kal-1</sub>::GFP]* IV; *mig-2(gm38gf)* X | Throughout |
| DV2224 | *otIs33[P<sub>kal-1</sub>::GFP]* IV; *rgl-1(ok1921Δ)* *mig-2(gm38gf)* X | Fig. 1F, S1F |
| DV2883 | *rgl-1(gk275304rf)* X | Starting reagent |
| DV2884 | *rgl-1(gk275305\*)* X | Starting reagent |
| DV2926 | *ral-1(gk628801rf)* III; *otIs33[P<sub>kal-1</sub>::GFP]* IV | Fig. S2C |
| DV2927 | *ral-1(gk628801rf)* III; *otIs33[P<sub>kal-1</sub>::GFP]* IV; *mig-2(gm38gf)* X | Fig. 3C |
| DV2942 | *ral-1(gk628801rf)* III | Starting reagent |
| NG2484 | *vab-8(gm84)* V | Starting reagent |
| OH904 | *otIs33[P<sub>kal-1</sub>::GFP]* IV | Throughout |
| CB315 | *unc-34(e315)* V | Starting reagent |
| DV2211 | *otIs33[P<sub>kal-1</sub>::GFP]* IV; *rgl-1(tm2255Δ)* *mig-2(gm38gf)* X | Fig. 1E, S1K |
| DV3565 | *otIs33[P<sub>kal-1</sub>::GFP]* IV; *vab-8(gm84)* V | Fig. 1I |
| DV3566 | *otIs33[P<sub>kal-1</sub>::GFP]* IV; *unc-34(e315)* V | Fig. S1C |
| DV3567 | *otIs33[P<sub>kal-1</sub>::GFP]* IV; *vab-8(e1017)* V | Fig. 1K |
| DV3568 | *otIs33[P<sub>kal-1</sub>::GFP]* IV; *rgl-1(ok1921Δ)* X | Fig. S1B |
| DV3577 | *otIs33[P<sub>kal-1</sub>::GFP]* IV; *rgl-1(tm2255Δ)* X | Throughout |
| DV3583 | *otIs33[P<sub>kal-1</sub>::GFP]* IV; *unc-34(e315)* V; *rgl-1(ok1921Δ)* X | Fig. S1E |
| DV3584 | *otIs33[P<sub>kal-1</sub>::GFP]* IV; *vab-8(gm84)* V; *rgl-1(ok1921Δ)* X | Fig. 1J |
| DV3615 | *otIs33[P<sub>kal-1</sub>::GFP]* IV; *rgl-1(gk275304rf)* X | Fig. S2B |
| DV3617 | *otIs33[P<sub>kal-1</sub>::GFP]* IV; *rgl-1(gk275305\*)* X | Fig. S2A |
| DV3619 | *otIs33[P<sub>kal-1</sub>::GFP]* IV; *unc-34(e315)* V; *rgl-1(tm2255Δ)* X | Fig. S1D |
| DV3621 | *otIs33[P<sub>kal-1</sub>::GFP]* IV; *rgl-1(gk275305\*)* *mig-2(gm38gf)* X | Fig. 3A |
| DV3238 | *ral-1(re160gf[mKate-2<sup>3</sup>xFlag::ral-1(G26V)])* III | Starting reagent |
| DV3402 | *ral-1(re218[mKate-2<sup>3</sup>xFlag::ral-1])* III | Starting reagent |
| DV3378 | *rheb-1(tm4642)* / *qC1 [dpy-19(e1259ts) glp-1(q339) nIs189[P<sub>myo-2</sub>::GFP]]* III | *ral-1* balancer |
| DV3631 | *otIs33[P<sub>kal-1</sub>::GFP]* IV; *rgl-1(gk275304rf)* *mig-2(gm38gf)* X | Fig. 3B |
| DV3633 | *ral-1(re218[mKate-2<sup>3</sup>xFlag::ral-1])* III; *otIs33[P<sub>kal-1</sub>::GFP]* IV; *mig-2(gm38gf)* X | Fig. 4A, 4B, S3B |
| DV3647 | *ral-1(re160gf[mKate-2<sup>3</sup>xFlag::ral-1(G26V)])* III; *otIs33[P<sub>kal-1</sub>::GFP]* IV; *mig-2(gm38gf)* X | Fig. 4A, 4B, S3B |
| LE732 | *lqls27[P<sub>ceh-23</sub>::GFP + lin-15]* IV; *lin-15B&lin-15A(n765)* X | Fig. S1H |
| DV3663 | *ral-1(re218[mKate-2<sup>3</sup>xFlag::ral-1])* III; *otIs33[P<sub>kal-1</sub>::GFP]* IV | Fig. S3A |
| DV2212 | *rgl-1(tm2255Δ)* *mig-2(gm38gf)* X | Starting reagent, Fig. 2A |
| MT2124 | *let-60(n1046gf)* IV | Starting reagent |
| DV3672 | *reEx256[P<sub>ceh-23_L</sub>::GFP + P<sub>myo-2</sub>::mCherry]* | Results, GFP silencing |
| CZ13799 | *juls76[P<sub>unc-25</sub>::GFP + lin-15]* II | Fig. 2B, 2C |
| DV3681 | *ral-1(re160gf[mKate-2<sup>3</sup>xFlag::ral-1(G26V)])* III; *otIs33[P<sub>kal-1</sub>::GFP]* IV | Fig. S3A |
| PS436 | *let-60(sy93dn)* IV | Starting reagent |
| DV2198 | *otIs33[P<sub>kal-1</sub>::GFP] IV; rgl-1(tm2255Δ) mig-2(gm103gf) X* | Fig. 1H |
| DV3690 | *lqls27[P<sub>ceh-23</sub>::GFP + lin-15] IV; rgl-1(tm2255Δ) mig-2(gm38gf) X* | Fig. S1I |
| DV3731 | *juls76[P<sub>unc-25</sub>::GFP + lin-15] II; mig-2(gm38gf) X* | Fig. 2B, 2D |
| DV3732 | *juls76[P<sub>unc-25</sub>::GFP + lin-15] II; rgl-1(tm2255Δ) mig-2(gm38gf) X* | Fig. 2B, 2E |
| MT20108 | *dpy-17(e164) unc-32(e189) / qC1[dpy-19(e1259ts) glp-1(q339) nls281[P<sub>myo-2</sub>::RFP]] III* | *ral-1* balancer |
| DV3734 | *otIs33[P<sub>kal-1</sub>::GFP] let-60(n1046gf) IV* | Fig. S3C |
| NG103 | *mig-2(gm103gf) X* | Starting reagent, Fig. 2A |
| NG2475 | *mig-2(gm38gf) X* | Starting reagent, Fig. 2A |
| DV2197 | *rgl-1(tm2255Δ) mig-2(gm103gf) X* | Fig. 2A |
| DA711 | *unc-10(e102) dpy-6(e14) X* |  |
| DV3736 | *otIs33[P<sub>kal-1</sub>::GFP] IV; vab-8(e1017) V; rgl-1(ok1921Δ) X* | Fig. 1L |
| DV3742 | *lqls27[P<sub>ceh-23</sub>::GFP + lin-15] IV; mig-2(gm38gf) X* | Fig. S1H, S1I |
| DV3152 | *rap-1(tm861Δ) IV* | Starting reagent |
| TU2589 | *dpy-20(e1282) IV; uls25[P<sub>mec-18</sub>::GFP + dpy-20] X* |  |
| JK2958 | *dpy-11(e224) unc-42(e270) V / nT1[qIs51[P<sub>myo-2</sub>::GFP + P<sub>pes-10</sub>::GFP + P<sub>F22B7.9</sub>::GFP]] IV, V* | *lin-45*, *let-60*, and *rgl-1* balancer |
| DV3755 | *otIs33[P<sub>kal-1</sub>::GFP] let-60(n1046gf) IV; mig-2(gm38gf) X* | Fig. 4C, S3D |
| DV2689 | *sec-5(pk2357) / mIn1[dpy-10(e128) mls14{P<sub>myo-2</sub>::GFP}] II* | Starting reagent |
| DV2690 | *rlbp-1(tm3665Δ) I* | Starting reagent |
| DV3202 | *exoc-8(ok2523Δ) I* | Starting reagent |
| DV3792 | *rlbp-1(tm3665Δ) I; otIs33[P<sub>kal-1</sub>::GFP] IV* | Fig. S3J |
| DV3793 | *exoc-8(ok2523Δ) I; otIs33[P<sub>kal-1</sub>::GFP] IV* | Fig. S3I |
| DV3795 | *rlbp-1(tm3665Δ) I; otIs33[P<sub>kal-1</sub>::GFP] IV; mig-2(gm38gf) X* | Fig. 4I |
| DV3796 | *exoc-8(ok2523Δ) I; otIs33[P<sub>kal-1</sub>::GFP] IV; mig-2(gm38gf) X* | Fig. 4H |
| MT4954.5 | *dpy-9(e12) ced-2(e1752) unc-33(e204) IV* |  |
| DR105 | *unc-17(e245) dpy-20(e1282) IV* |  |
| DR281 | *unc-31(e169) dpy-4(e1166) IV* |  |
| DV3811 | *mIn1[dpy-10(e128) mls14{P<sub>myo-2</sub>::GFP}] II; otIs33[P<sub>kal-1</sub>::GFP] IV; mig-2(gm38gf) X* | Starting reagent |
| DV3820 | *reSi8[P<sub>ceh-23_L</sub>::mKate-2::ral-1(+)] I; ral-1(gk628801rf) III; otIs33[P<sub>kal-1</sub>::GFP] IV* | Fig. S4 A-D |
| DV3821 | *reSi9[P<sub>ceh-23_L</sub>::mKate-2::ral-1(+)] I; ral-1(gk628801rf) III; otIs33[P<sub>kal-1</sub>::GFP] IV* | Fig. S4 E |
| DV3834 | *otIs33[P<sub>kal-1</sub>::GFP] rap-1(tm861Δ) IV; mig-2(gm38gf) X* | Fig. 4E |
| DV3487 | *lin-45(ku112) let-60(n1046gf) IV* | Starting reagent |
| DV3849 | *reSi14[P<sub>ceh-23_L</sub>::mKate-2::rgl-1(+)] I; otIs33[P<sub>kal-1</sub>::GFP] IV; rgl-1(tm2255Δ) X* | Fig. 5A-D |
| DV3850 | *reSi15[P<sub>ceh-23_L</sub>::mKate-2::rgl-1(+)] I; otIs33[P<sub>kal-1</sub>::GFP] IV; rgl-1(tm2255Δ) X* | Fig. 5E |
| DV3874 | *sec-5(pk2357) / mln1[dpy-10(e128) mls14{P<sub>myo-2</sub>::GFP}] II;*<br>*otIs33[P<sub>kal-1</sub>::GFP] IV; mig-2(gm38gf) X* | Fig. 4G |
| DV3875 | *sec-5(pk2357) / mln1[dpy-20(e128) mls14{P<sub>myo-2</sub>::GFP}] II;*<br>*otIs33[P<sub>kal-1</sub>::GFP] IV* | Fig. S3H |

**Supplementary Table 2:**
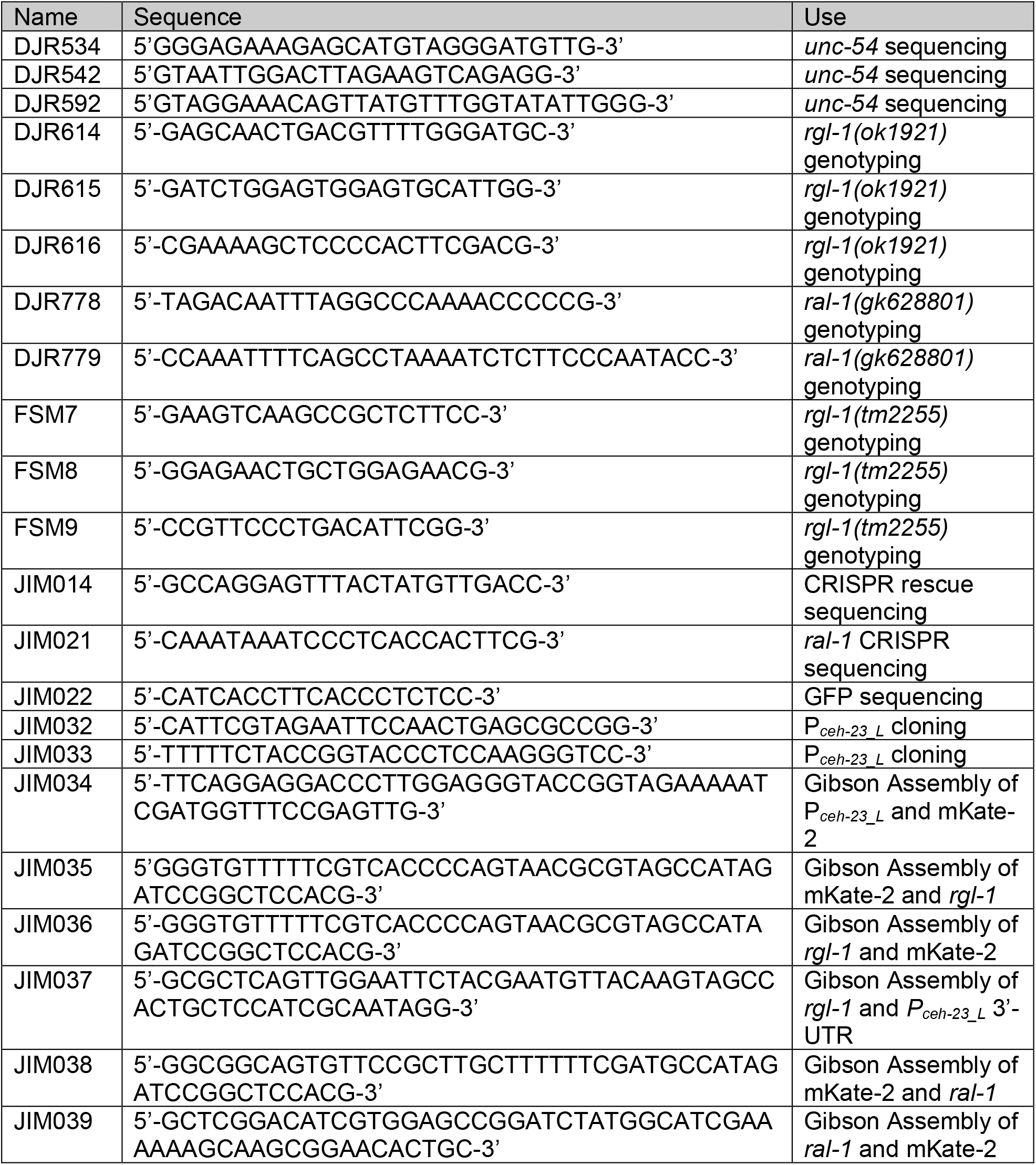
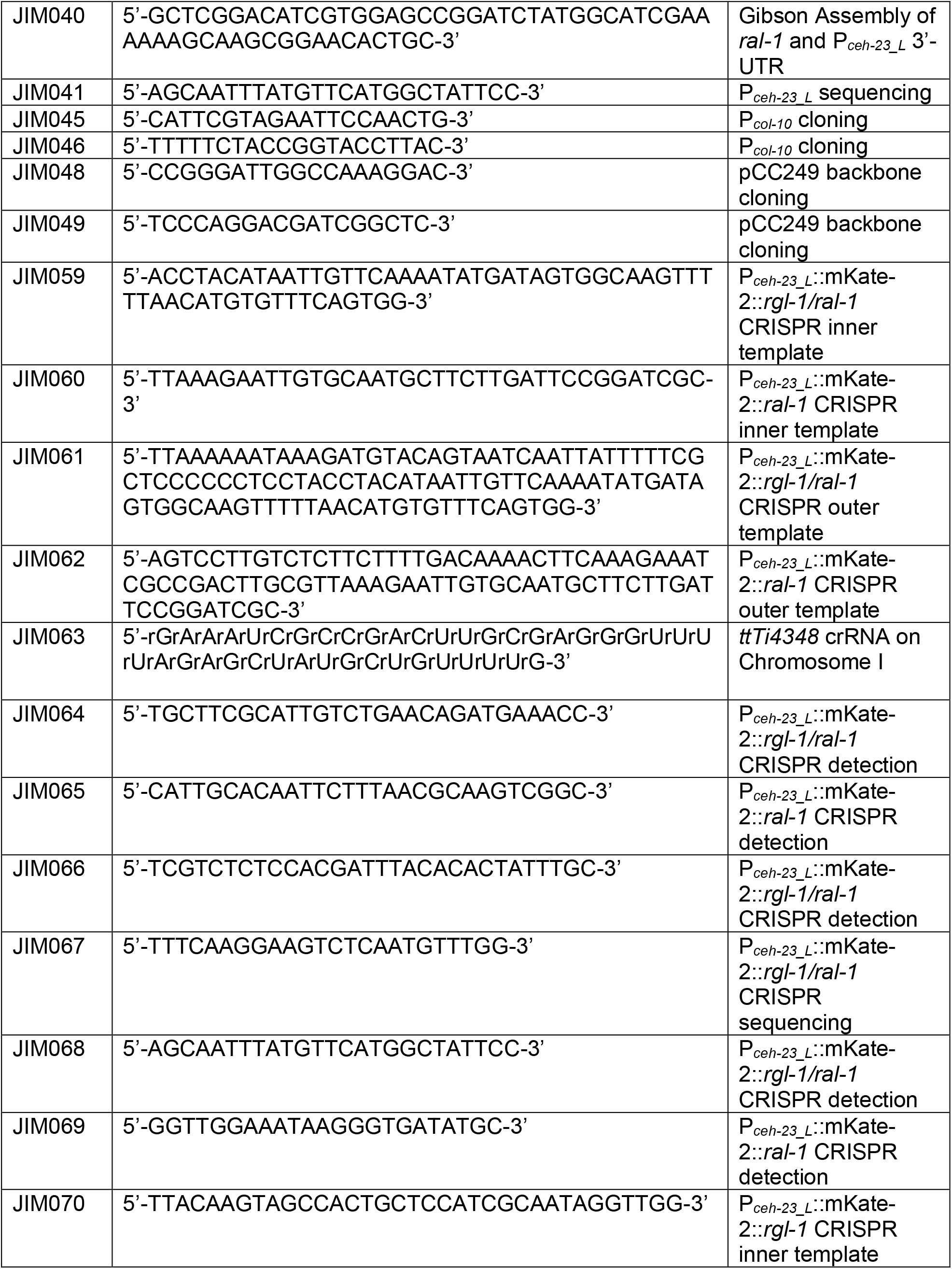
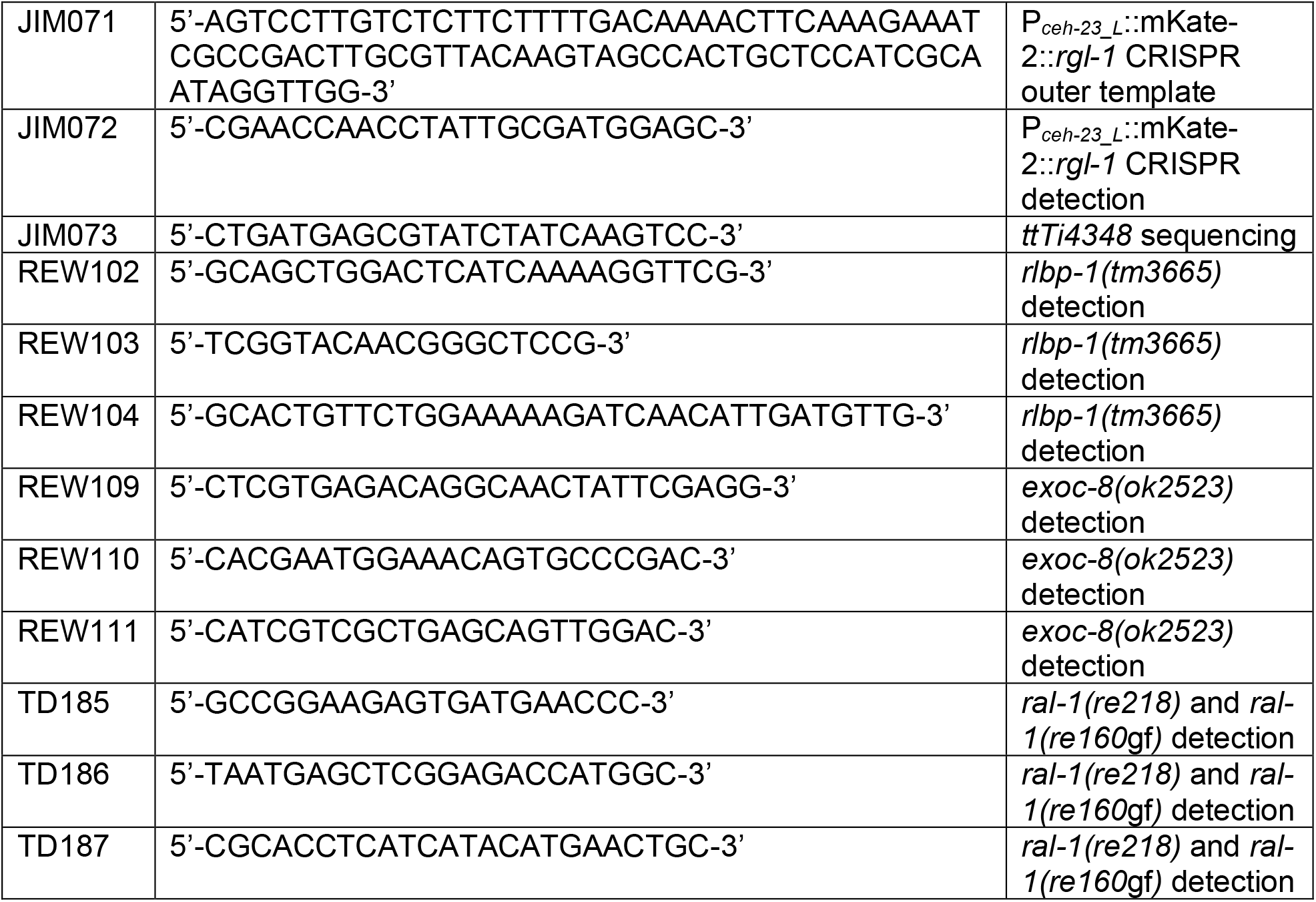
Primers.

